# Retro-cue alerting effects on conscious perception across different levels of target visibility

**DOI:** 10.64898/2026.03.13.711604

**Authors:** Pablo Rodríguez-San Esteban, Mariagrazia Capizzi, José A. González-López, Ana B. Chica

## Abstract

Can alerting retro-cues improve the conscious perception of stimuli that would otherwise remain unseen? While pre-stimulus alerting is known to facilitate conscious access, the effects of retro-cues remain ambiguous due to methodological confounds in existing literature. Specifically, most studies finding retro-cue benefits have relied on spatial features (such as lateralized targets or cues), which confound alerting with spatial selection. Our design addresses this gap by employing central visual targets and non-lateralized auditory cues, thereby isolating the temporal boost of phasic alerting from spatial orienting. Across four experiments, participants reported the presence and orientation of a central Gabor patch presented at near-threshold (∼50% detection)) or higher visibility (∼75% detection) levels. An auditory alerting tone was presented prior to, simultaneously with, or after the Gabor, at various short and long stimulus onset asynchronies, with both short and long temporal ranges. Results consistently showed that pre-stimulus and simultaneous cues significantly enhanced conscious perception, increasing both seen rates and (in some experiments) perceptual sensitivity. Crucially, the presence of retro-cue benefits was observed at higher stimulus visibility, but not at near-threshold, suggesting a potential influence of target visibility on retro-cue efficacy. When targets were near-threshold, retro-cues did not reliably improve conscious perception, whereas tones presented 200 ms after target offset significantly increased the proportion of seen reports when target visibility was increased . These findings indicate that nonspatial auditory retrocues can modulate conscious report when the underlying sensory trace is sufficiently robust. More broadly, they show that the timing of conscious access can be influenced by late alerting signals even in the absence of spatial information, providing a behavioural platform to refine models of how alerting and evidence strength interact in conscious perception.

## 1 Introduction

Attention and conscious perception are closely related (Dehaene et al., 2006; Posner, 1994), with some authors defending that attention serves as a gateway to consciousness (Chica and Bartolomeo, 2012; Dehaene, 2001; Dehaene et al., 2006; Posner, 1994). This fundamental relationship between attention and consciousness has been examined with respect to each of the three attentional networks proposed by Posner and Petersen (2012, 1994, 1990), exploring their specific links to consciousness. The orienting system allows selecting a certain location in space, which in turn makes the information presented at that location more accessible and enhancing the conscious perception of stimuli (Chica et al., 2012). The executive control network is responsible for conflict resolution, error detection, planning and decision making, particularly in novel situations (Fan et al., 2005; Fan et al., 2002; Posner and Dehaene, 1994). When this system is highly activated, for instance, during a Stroop task, the conscious perception of stimuli can be impaired, suggesting that these processes share underlying neural circuits (Colás et al., 2017, 2018; Martín-Signes et al., 2019, 2021). Lastly, the alerting network is critical for maintaining a state of activation and vigilance, enabling individuals to react to stimuli or changes in their environment (Posner and Petersen, 1990). Alerting can be either tonic (intrinsic), which refers to a sustained state of alertness, or phasic (extrinsic), which refers to a transient increase in alertness in response to a specific stimulus (Posner and Petersen, 1990; Sturm and Willmes, 2001). Phasic auditory alerting has been consistently shown to enhance both objective and subjective measures of conscious perception (Botta et al., 2014; Chica et al., 2016; Kusnir et al., 2011; A. Petersen et al., 2017; Poth, 2025). This enhancement occurs by increasing visual processing speed and lowering the threshold for conscious perception, with the magnitude of this effect scaling with cue intensity (A. Petersen et al., 2017). A recent study by Dupont et al. (2026) employed a purely accuracy-based paradigm combined with computational modelling to demonstrate that phasic alerting acts as a global sensory amplifier, increasing visual processing speed while remaining independent of tonic arousal level. Crucially, by employing non-aging foreperiod distributions to minimize temporal expectancy, their work reinforces the idea that these alerting benefits reflect a transient boost in perceptual readiness rather than a sustained shift in baseline activation or strategic anticipation (Dupont et al., 2026). Furthermore, a direct link has been established between cue-induced desynchronization of the occipital alpha rhythm and improved visual discrimination, with these beneficial effects being mediated by fronto-parietal networks that provide top-down amplification of visual processing (Chica et al., 2016; Feng et al., 2017). This suggests that auditory alerting cues can modulate the neural mechanisms underlying conscious perception, enhancing the processing of visual stimuli and facilitating their access to conscious awareness.

However, the enhancing effects of phasic alerting have primarily been investigated when the alerting cues are presented prior to the target stimulus (Chica et al., 2016; Kusnir et al., 2011; A. Petersen et al., 2017). When the tone is presented after the target stimulus (which is known as retro-cue), its influence on conscious perception is less clear. Retro-perception refers to the phenomenon where attracting attention to a stimulus location after its disappearance can retrospectively trigger its conscious perception, even when it would otherwise have gone unseen (Sergent et al., 2013). This effect is characterized by a discrete, all-or-none transition in conscious perception, meaning retro-cues increase the likelihood of perceiving a stimulus (reducing guess trials) rather than simply improving the precision or fidelity of an already conscious representation (Thibault et al., 2016). A key implication is that, according to late and global approaches, conscious perception is not strictly time-locked to the initial sensory input but it can be desynchronized from it, becoming conscious at a different moment than the initial timing of the stimulus (Sergent, 2018). This suggests a temporal flexibility in conscious perception, allowing an event occurring hundreds of milliseconds after the stimulus to determine its conscious fate (Sergent et al., 2013).

Historically, retro-cues were thought to improve performance by facilitating the transfer of information from a high capacity, short-lived sensory memory (like iconic memory) to a more stable, but limited, working memory system, without affecting perception itself (Sergent et al., 2013). However, more recent research, particularly on retro-perception, directly challenges this assumption by proposing that retro-cues can directly influence whether a stimulus is consciously perceived, not just how well it is remembered if it was already conscious (Garnier-Allain et al., 2023). This highlights a perception-memory continuum, suggesting that perception is not fully determined at the initial sensory stage but remains malleable and open to influences even after stimulus offset (Szaszkó et al., 2024). Retro-cue paradigms have also been widely used to investigate how information is recovered or prioritized from working memory (Griffin and Nobre, 2003; Guo et al., 2024; Landman et al., 2003; Lin et al., 2021; Loaiza et al., 2024; Schneider et al., 2015; Souza and Oberauer, 2016; Wildegger et al., 2016; Zerr et al., 2021). In these studies, a retro-cue presented after the initial encoding phase directs attention to a specific item or location in memory, allowing researchers to probe the mechanisms by which attention can enhance the accessibility or fidelity of stored representations. This body of work has demonstrated that retro-cues can improve recall accuracy and reduce interference, highlighting the dynamic nature of working memory and the role of attention in modulating its contents.

While some studies suggest that retro-cues can enhance the perception of past stimuli, the evidence for such enhancement is significantly richer when visual cues are employed (Garnier-Allain et al., 2023; Gressmann and Janczyk, 2016; Sergent, 2018; Szaszkó et al., 2024; Thibault et al., 2016; Xia et al., 2016; Zerr et al., 2021). However, a critical characteristic of these paradigms is that they typically involve lateralization of the stimulus, the cues, or both. For example, even in the auditory domain, retro-cueing benefits often involve selecting between streams (e.g., left vs. right ear) or identifying low-level features like pitch from a multi-item array. Consequently, these studies do not only examine a pure temporal alerting effect but also a spatial or feature-based selection effect (Abdi, 2010; Astle et al., 2012; Gressmann and Janczyk, 2016; Griffin and Nobre, 2003; Janczyk and Reuss, 2016; Rimsky-Robert et al., 2019; Schneider et al., 2016; Sergent, 2018; Sergent et al., 2013; Szaszkó et al., 2024; Thibault et al., 2016; Xia et al., 2016).

To the best of our knowledge, all previous studies that have reported significant behavioral retro-cue benefits have incorporated some form of spatial information (Griffin and Nobre, 2003; Landman et al., 2003; Souza and Oberauer, 2016). This is typically achieved either by lateralizing the target stimulus or by using cues that contain explicit spatial information, such as presenting an auditory tone in only one ear or from a specific loudspeaker (Rimsky-Robert et al., 2019; Rummukainen and Mendonça, 2016; Sergent et al., 2013; Xia et al., 2016). Furthermore, the temporal window to observe retro-cueing effects appears to be highly limited; for instance, Rimsky-Robert et al. (2019) found significant retro-active improvements in detection sensitivity and reaction times at a +150 ms intervals, but these benefits were markedly reduced or absent at a longer delay of +450 ms. This indicates that the capacity of a retro-cue to modulate perception is consistent with the brief persistence of sensory representations, which decay rapidly after stimulus offset.

Crucially, there are no studies to date that have found significant retro-cue benefits using purely non-spatial alerting tones for the retrospective triggering of conscious perception, particularly in paradigms involving near-threshold Gabor patches (Keefe et al., 2021; Paladini et al., 2018). This systematic reliance on spatial features can confound the interpretation of existing results, making it difficult to disentangle whether observed improvements in perception are due to enhanced alertness, improved spatial selection, or an interaction of both (Chun et al., 2011; Posner and Petersen, 1990; Weinbach and Henik, 2012a). As a result, the effects attributed to retro-cues in the literature may not purely reflect the temporal dynamics of alerting but could be fundamentally driven by the additional benefits of spatial attention.

Therefore, a central question in consciousness research concerns the temporal relationship between sensory input and conscious experience: is conscious perception fixed at the moment of stimulus onset, or can it be triggered later, decoupled from the initial sensory event? Demonstrations of retro-perception and retrospective cueing in both vision and audition suggest that, under some conditions, awareness can be triggered by events occurring hundreds of milliseconds after stimulus offset, indicating a surprising temporal flexibility of conscious access (Sergent et al., 2013; Thibault et al., 2016). In this framework, alerting retro-cues provide a powerful way to probe whether late, non-specific signals can promote existing sensory traces into awareness, thereby constraining theories that differ in how tightly they tie conscious access to initial encoding versus later global amplification (Ciupiska et al., 2024; Dehaene et al., 2006; Sergent, 2018). However, most demonstrations of retro-perception have relied on spatially informative cues or lateralized stimuli, making it difficult to determine whether the critical factor is temporal alerting per se or the well-known benefits of spatial orienting (Griffin and Nobre, 2003; Sergent et al., 2013; Souza and Oberauer, 2016). Isolating the pure temporal component of retrospective attention is therefore essential for understanding how alerting contributes to conscious access and for providing a cleaner behavioural platform to test competing models of consciousness (Ciupiska et al., 2024; Lawrence and Klein, 2013).

This study aimed to address these existing ambiguities by systematically exploring the effects of auditory retro-cues on the conscious perception of a central target stimulus. To achieve this goal, we designed an experimental paradigm that allowed manipulating the timing of an auditory alerting cue relative to the onset of a near-threshold Gabor (see Rodríguez-San Esteban et al., 2025). A key aspect of our experimental design was the use of central visual stimuli and non-lateralized auditory cues. Unlike many previous studies that involve lateralization of stimuli or cues (which can introduce validity or congruency effects) (Rimsky-Robert et al., 2019; Sergent, 2018; Sergent et al., 2011, 2013; Thibault et al., 2016), our approach focuses on a central alerting mechanism without such spatial confounds. This allows for a clearer examination of how the temporal relationship between a non-spatial auditory cue and a central visual target influences conscious perception.

## 2 General methods

### 2.1 Overview of the experimental series

While the core experimental task and stimuli remained consistent across the four experiments presented in this paper, each experiment introduced specific modifications to test the alerting effect under different conditions of temporal predictability and cue-to-target asynchrony. Below, we provide a general overview of the main manipulations employed in each experiment (see Table 1).

**Table 1.** Summary of experimental parameters used across the four experiments.

| Experiment | Fixation duration | SOAs | Target visibility (% seen trials) |
| --- | --- | --- | --- |
| Experiment 1 | Fixed (1000 ms) | -200, -100, 0, +100, +200 | ~50% |
| Experiment 2 | Variable (1000–1500 ms) | -200, -100, 0, +100, +200 | ~50% |
| Experiment 3 | Variable (1000–1500 ms) | -400, -200, 0, +200, +400 | ~50% |
| Experiment 4 | Variable (1000–1500 ms) | -400, -200, 0, +200, +400 | ~75% |

### 2.2 Participants

From the complete sample of each experiment, outliers were removed based on the average proportion of seen targets and false alarms (FA) in the detection task. Outliers were defined as those participants whose average proportion of seen targets and/or FA was above or below 2 standard deviations from the mean. This criterion was established prior to any data inspection, and the same data exclusion procedure was applied to all four experiments.

Although an *a priori* power analysis was not conducted before data collection, we used the data from Experiment 1 as a “pilot” or baseline study to evaluate the sensitivity and robustness of our project-wide sample size (*N* = 35) across the entire experimental series (Lakens, 2022). This post-hoc design analysis (Gelman and Carlin, 2014) evaluates the properties of the experimental design by using the specific variance components and noise estimated from Experiment 1 to establish its statistical resolution based on empirically derived parameters (Kruschke, 2015), and was run after the experimental series was completed. Our implementation follows the logic of standardized simulation-based power analysis frameworks such as the *simr* package for R (Green and MacLeod, 2016), adapted for the Python environment using custom Monte Carlo simulations based on the *pymer4* library. First, a simulation-based sensitivity analysis (Brysbaert and Stevens, 2018) showed that at N = 35, we maintained >80% power (estimated via 100 simulations) to detect alerting effects larger than 45 ms in reaction times (RTs) and 5% in accuracy (ACC) or proportion of seen. Second, a non-parametric subsampling analysis (Green and MacLeod, 2016) confirmed that our core alerting effects reached the 80% power threshold at sample sizes of *N* = 20 for proportion of seen, *N* = 30 for ACC, and *N* = 25 for RTs. These analyses collectively demonstrate that our project-wide sample of *N* = 35 provided high statistical sensitivity and robust detection of the reported experimental effects. All following experiments maintained the same sample size of *N* = 35 to ensure consistency and comparability.

In **Experiment 1**, 40 participants were recruited (27 females, mean age = 21.55, SD = 3.41). Two participants were removed from the sample due to failure in completing the titration procedure, and three were discarded due to outlier values in the average proportion of seen and/or FA. In **Experiment 2**, the study sample included 42 volunteers (26 females, mean age 22.71 years, SD = 2.8). Some participants had to be removed from the sample due to technical issues (1 participant), not being able to successfully complete the titration procedure (3 participants) and not reaching higher-than-chance accuracy in the orientation discrimination task (3 participants). In **Experiment 3**, the initial sample included 42 participants (30 females, mean age 21.5, SD = 3.08). Some subjects had to be removed from the sample due to failure in completing the titration procedure (3 participants) or having outlier values in the average proportion of Seen and/or FA (4 participants). In **Experiment 4**, 45 participants were recruited (35 females mean age = 20.26, SD = 1.89). Some participants (5) were removed from the sample due to failure in completing the titration procedure, and others were discarded due to outlier values in the average proportion of Seen and/or FA (5 participants). The final sample included in each experiment and used for the analyses was of 35 participants.

All participants had normal or corrected-to-normal vision, and no prior experience with the task. No participant had a history of major medical, neurological, or psychiatric disorders. Participants received a monetary compensation of 5/h for their participation and were recruited using the SONA experiment management system. Signed informed consent was collected prior to their inclusion in the study. articipants were informed about their right to withdraw from the experiment at any time. The University of Granadas Research Ethics Committee approved the experiment (1862/CEIH/2020), which was carried out in accordance with the Code of Ethics of the World Medical Association (Declaration of Helsinki) for experiments involving humans (last update: Brazil, 2013).

### 2.3 Apparatus and stimuli

PsychoPy version 2023.2.3 (Peirce et al., 2019) was used to present stimuli and collect task responses. Participants were seated at an approximate distance of 56 cm from the computer screen (a 24” monitor, BenQ XL2411T, 1920x1080 pixels, with a refresh rate of 60 Hz), and an adjustable headrest was used to stabilize head position and ensure consistent distance.

Trials started with a fixation display, consisting of a plus sign (0.71^°^x 0.71^°^) that was displayed in the center of the screen. The target was a Gabor stimulus generated using PsychoPy built-in functions (phase = 0.25 cycles, spatial frequency = 2.025, 3.58^°^diameter), and a 500Hz tone was also generated using PsychoPy, presented at 70dB through headphones. The mask consisted of a checkerboard pattern (3.06^°^ diameter), presented largely above threshold contrast. In the tilt orientation response screen, two Gabor probes, identical to the target stimulus, were presented 1^°^above and below the fixation cross. Finally, the words “Visto” (“Seen”, in Spanish; 3.06^°^ x 1^°^) and “No visto”(“Unseen”, in Spanish; 5.31^°^x 1^°^) were displayed 1^°^above and below the fixation cross.

### 2.4 Task and procedure

Each experiment started with an initial titration procedure, conducted to adjust Gabor contrast to achieve either 50% (in Experiments 1, 2, and 3) or 75% (in Experiment 4) proportion of seen Gabors during the experimental blocks (see description of this procedure below, and see Table 1 for a general overview of the main manipulations employed in each experiment). Then, participants performed a total of 400 trials (300 Gabor present and 100 Gabor absent, randomly presented). During these trials, the contrast for the present stimuli was set to that obtained during the titration procedure. The total duration of the experimental session was approximately 70 minutes.

Trials in all blocks had a similar design (see Figure 1). They started with a fixation screen, with a fixed duration of 1000 ms (Experiment 1), or variable duration ranging between 1000 ms and 1500 ms (Experiments 2 and 3). After this initial fixation, the Gabor stimulus was displayed at fixation during 50 ms, tilted clockwise or counterclockwise (with equal probability). In catch trials, no Gabor stimulus was presented. An alerting tone could be present, equally distributed between six stimulus onset asynchrony (SOA) conditions. One was always a “no tone” condition, with no sound, and the others were experiment-dependent: in Experiment 1 and Experiment 2, the SOAs were (respective to the Gabor onset) -200 ms, -100 ms, 0 ms, +100 ms, and +200 ms, while in Experiment 3 the SOAs were -400 ms, -200 ms, 0 ms, +200 ms, and +400 ms. his tone was described as task-irrelevant to the participants, and could be displayed in both target present and target absent trials. After 300 ms, the mask was presented for 100 ms, followed by an inter-stimulus-interval (ISI) of 1950 ms, in which the fixation cross was displayed. Participants were then required to report the orientation of the lines composing the Gabor. To do this, they were presented with a clockwise-oriented Gabor, and a counter-clockwise oriented Gabor, above and below the fixation cross, with the location of these stimuli being randomized across trials to avoid response preparation. This screen was presented until response. Participants responded with their right hand, pressing either the “k” key associated with the stimulus above the fixation point or the “m” key for the one presented below the fixation point (right hand response). Participants were asked to randomly press one of the two keys if no Gabor was perceived. Subsequently, participants were required to report the presence or absence of the Gabor, with a screen displaying the words “Seen” (“Visto”, in Spanish) and “Unseen” (“No visto”, in Spanish) above and below the fixation point, with the location of these words being also randomized across trials to avoid response preparation. This screen was displayed until a response was detected. Participants responded with their left hand, using the keys “d” for the response above the fixation point and “c” for the one below the fixation point. Finally, an inter-trial interval with a duration of between 1500 ms and 2000 ms was presented with the fixation cross displayed on the centre of the screen.

**Figure 1.**
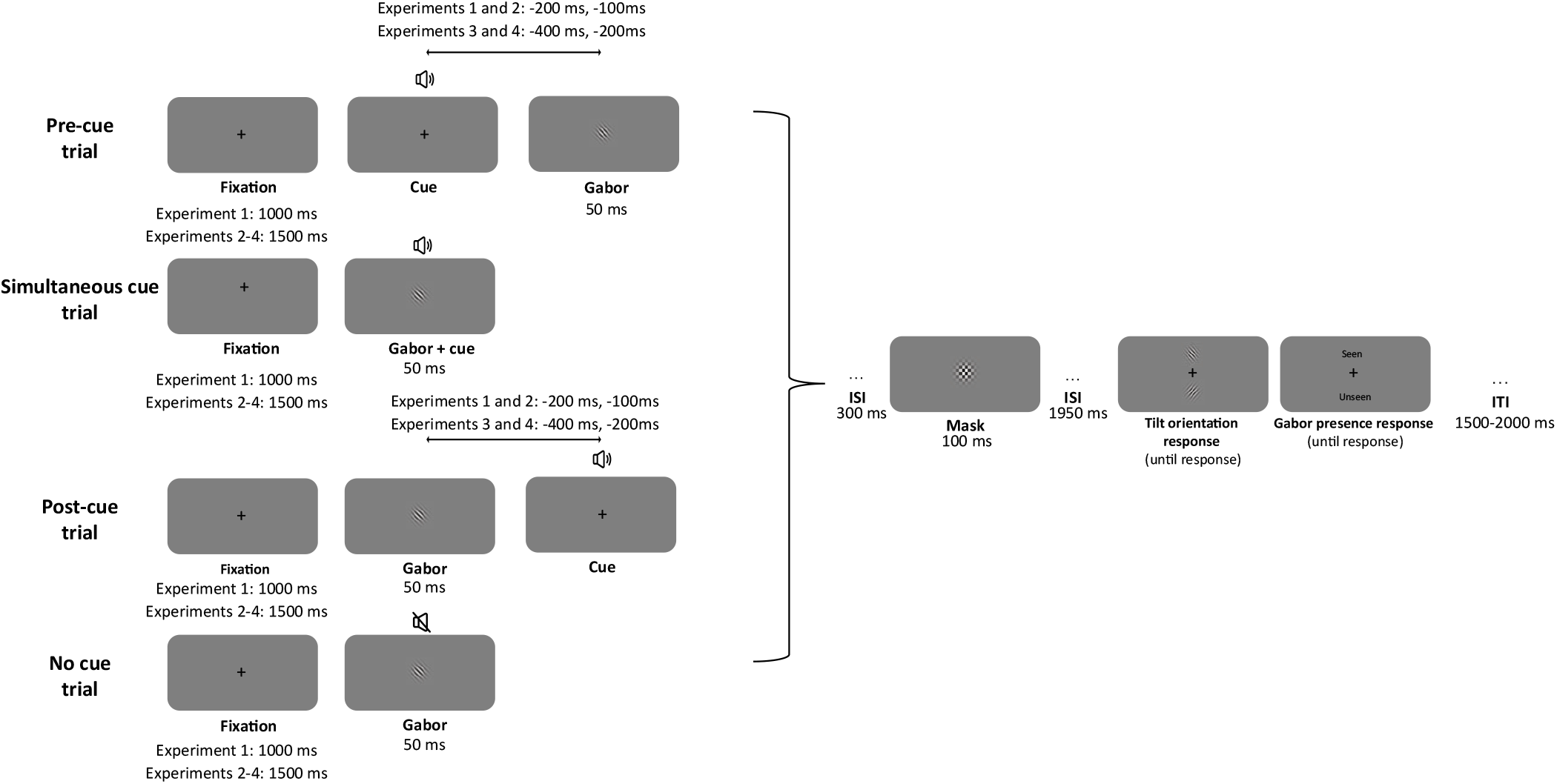
Schematic representation of the experimental task, showing the different conditions: pre-cue (tone presented before the target stimulus), simultaneous (both presented at the same time), post-cue (tone presented after the target stimulus), or no tone trials. Each trial consisted of a fixation period, presentation of the Gabor stimulus (or catch trials, with no Gabor), an alerting tone at varying SOAs depending on the experiment, a mask, and subsequent response screens for orientation discrimination and subjective awareness.

During the initial titration block, which was used to adjust the percentage of seen targets individually, two consecutive QUEST (Watson and Pelli, 1983) staircases (30 trials each) were conducted as implemented in PsychoPy. The first staircase started with a supra-threshold Gabor, and the second staircase started with the last contrast value of the previous QUEST. In both of these staircases only target present trials were performed. Then, an additional 60 trials block was run to allow for a finer modulation of the target contrast. The initial contrast for this block was the average contrast of the last 20 trials in the second QUEST staircase, and was titrated on a trial-by-trial basis following a one-up and one-down strategy: if participants reported seeing a stimulus, the contrast was decreased by 0.001 units, and if they reported not seeing the stimulus, the contrast was increased by 0.001 units. It is important to note that in this block there were catch trials (33.33%). At the end of this block, the proportion of seen stimuli and FA was computed, and a decision was made: if the proportion of seen targets ranged between 45% and 55% (for the first three experiments), or between 70% and 80% (in Experiment 4), and the proportion of FA was lower than 30%, the titration procedure was finished and the experimental trials started. If any of the conditions was not met, the practice block was repeated up until a maximum of three times. If, after the third block, the percentage of seen targets did not stabilize and/or the proportion of FA was too high, the participant was disqualified to complete the session.

The PsychoPy task files, as well as the behavioural data, can be found at the following repository: https://doi.org/10.5281/zenodo.18999229.

### 2.5 Statistical analyses

The code for all these analyses can be found here: https://anonymous.4open.science/r/RetroCuesAlerting

#### 2.5.1 Linear mixed-effects models

Linear mixed models (LMMs) have gained popularity as an alternative to repeated-measures ANOVAs in the field of experimental psychology (Magezi, 2015), mainly because this framework offers various notable advantages over the traditional approach (Lo and Andrews, 2015). First, LMMs enable the explicit specification of random effects such as participant-level variations, rather than pooling this variability within an error term. Secondly, these models are more flexible when it comes to handling missing data as compared to ANOVAs, enhancing the robustness of the analysis. Finally, LMMs are also more robust to violations of distributional assumptions than traditional ANOVAs (Schielzeth et al., 2020).

LMMs were conducted to analyze behavioral effects, using the pymer4 Python package (Jolly, 2018) to calculate significance applying Satterthwaite’s method to estimate degrees of freedom. In three different analyses, we included participants log-transformed RTs, accuracy, and the proportion of seen targets as the dependent variables. We then compared different model structures, in which the fixed effects included the SOA. Then, in the random structure, the model could either include or not a random slope for the trial within each participant. All possible model structures were fitted and then compared using both the Akaike Information Criterion (AIC) and Bayesian Information Criterion (BIC), as well as log-likelihood values. The model with the lowest AIC and BIC was selected as the best model for each analysis. These were the model structures selected for each analysis, including a random intercept for the participant and a random slope for the trial within each participant in the random structure:

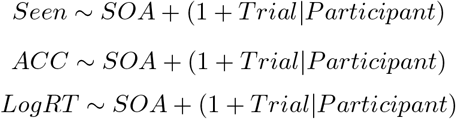

For the analyses of the proportion of seen responses and the ACC, the model assumed a binomial distribution (responses could only be either 0 or 1); for the RTs analysis, we first log-transformed the data to normalize its distribution. We then specified a model with a Gaussian family. To confirm this was the appropriate approach, we also tested alternative generalized linear mixed models (GLMMs) on the raw, non-transformed RTs using both Gamma and Inverse-Gaussian families. This model comparison revealed that these alternative GLMMs failed to converge to a stable solution for this dataset, as evidenced by unidentifiable parameters and zero-variance components for the random effects. Therefore, the stable and statistically valid Gaussian model was selected for all subsequent RTs analyses.

The significance of fixed effects was assessed by comparing the full model with a reduced model by means of likelihood-ratio tests (LRTs). The comparison was always run between nested models (i.e., the reduced model was the full model minus exactly one term, keeping the rest of the fixed-effect and random-effect structures the same). Then, both models were compared using the log-likelihood values, and the significance of the fixed effect was determined by the resulting Chi-squared metric and associated p-values. To qualify for any interactions, marginal estimates were computed and Bonferroni-corrected p-values were obtained for the post-hoc pairwise comparisons.

#### 2.5.2 Signal Detection Theory indices

Participant’s responses to the detection task were also analyzed using the Signal Detection Theory (SDT) (Abdi, 2007; Stanislaw and Todorov, 1999) framework. We obtained *d*^′^ and *β* values for each participant, which are measures of perceptual sensitivity and response criterion, respectively. *d*^′^ is a measure of the sensitivity to detect the Gabor stimulus, with higher values representing greater sensitivity. *β* indicates the response bias towards reporting the presence of the Gabor, with *β >* 1 representing a more conservative bias and *β <* 1 representing a liberal bias.

Catch trials (target-absent trials) were distributed across the same six SOA conditions as target-present trials. To avoid infinite values in *d*^′^ when FA rates were exactly 0, we applied a small log-linear style correction by replacing 0 with (0.5)/(N+1), where N was the mean number of catch trials per SOA, resulting in an adjusted FA rate of ≈0.028. Hit rates were not corrected because our inclusion criteria ensured that they remained within a predefined range and did not reach the extremes of 0 or 1.

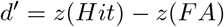

where *z* is the z-transform of the corresponding rates.

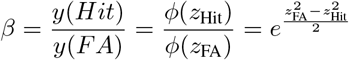

where:

- *y* is the ordinate (height) of the normal distribution at the computed z-value.
- *φ*(*z*) is the standard normal probability density function (PDF): 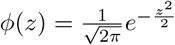
- *z*_Hit_ = Φ^−1^(1 − Hit Rate)
- *z*_FA_ = Φ^−1^(1 − False Alarm Rate)
- Φ^−1^ is the inverse standard normal CDF

Since indices from the SDT scores reported mean values and not trial-by-trial scores, the effect of SOA was analyzed by means of repeated-measures analysis of variances (ANOVAs), using the *Pingouin* Python package (Vallat, 2018). The assumptions of normality and homogeneity of variances were assessed using the Shapiro-Wilk test (Shapiro and Wilk, 1965) and Levene’s test (Levene, 1960), respectively. When violations of normality were detected, the non-parametric Friedman test (Friedman, 1937, 1940) was employed as an alternative to the repeated-measures ANOVA. Mauchly’s test was used to check for sphericity (Mauchly, 1940), and the Greenhouse-Geisser correction was applied to the degrees of freedom whenever the assumption of sphericity was violated (Abdi, 2010). When significant interactions were found, post-hoc pairwise comparisons (paired t-tests) were conducted using the Bonferroni correction for multiple comparisons. If the Friedman test was used, post-hoc analyses were performed using the Wilcoxon signed-rank test with Bonferroni correction.

#### 2.5.3 Linearity checks

To further explore how the alerting tone at different SOAs modulated the target detection, we examined whether the proportion of seen responses and the *d*^′^ values varied linearly or quadratically with SOA. For this purpose, we conducted polynomial contrasts within the LMMs framework. Specifically, we tested for linear and quadratic trends by including orthogonal polynomial terms for SOA as fixed effects in the models. The significance of these polynomial terms was assessed using Wald z-tests for the binomial GLMMs (for the proportion of seen responses) and t-tests (using Satterthwaite approximations for degrees of freedom) in the LMMs (for *d*^′^ values).

## 3 Results

To enhance the clarity of the manuscript and focus on the most critical findings, below we present the results for the Proportion of seen targets, ACC and SDT measures. All analyses regarding RTs are reported in the Supplementary Materials. Additionally, that section also includes the detailed Tables for pairwise post-hoc comparisons for Proportion of seen, ACC and SDT measures.

### 3.1 Experiment 1

As stated in the Methods section, in Experiment 1 the fixation screen duration was fixed to 1000 ms. This design provides predictable timing for stimulus appearance, allowing for a clearer assessment of the alerting effect under conditions of low temporal uncertainty and pure alerting effects (Fan et al., 2002). The SOAs for the auditory tone were −200 ms, −100 ms, 0 ms, +100 ms, and +200 ms relative to Gabor onset.

#### 3.1.1 Proportion seen

In the proportion of seen targets analysis, the paired-samples t-tests demonstrated a significant difference between Correct and Incorrect responses (*t*(35) = 58.086, *p* < .001, Cohen’s *d* = 1.102, BF_10_ > 1000), with correct trials having a higher proportion of seen targets than incorrect trials. Additionally, the overall proportion of FA (i.e., trials in which participants reported seeing the target when it was absent) in this experiments was .025 (SD = .026).

The LMMs revealed a significant effect of SOA (*χ*^2^(5) = 94.538, *p* < .001). Post-hoc pairwise comparisons (see Table S3) revealed a pattern of better performance (i.e., increased proportion of seen targets) when the alerting tone was presented prior to Gabor onset (Figure 2). All following post-hoc pairwise comparisons were Bonferroni-corrected.

**Figure 2.**
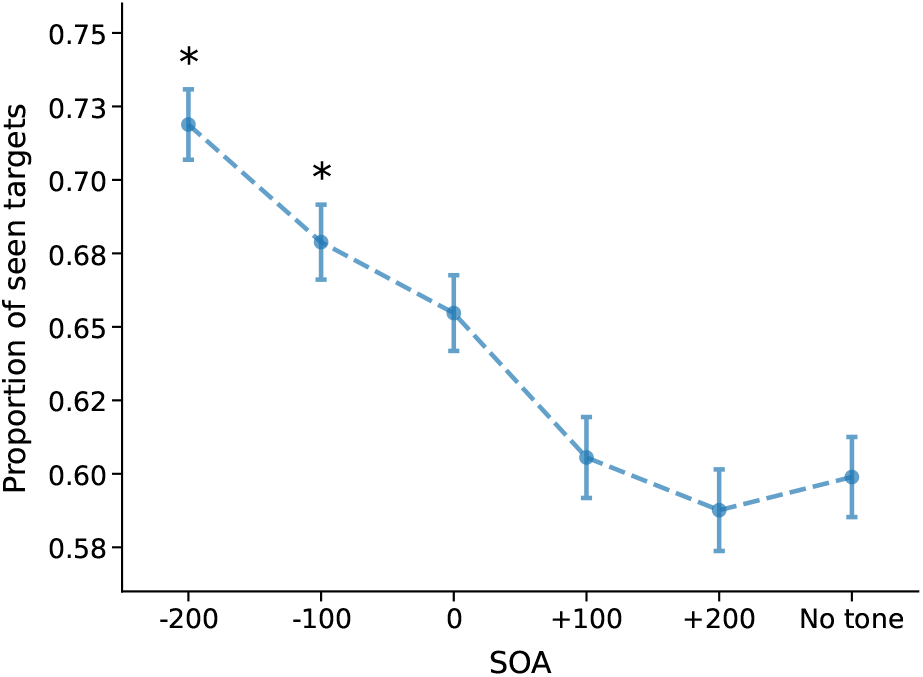
Proportion of seen targets for Experiment 1 as a function of SOA. Error bars represent standard errors. Asterisks indicate significant post-hoc comparisons against the No tone condition.

To further characterize the temporal dynamics of this effect, a GLMM was fitted with orthogonal polynomial contrasts for the fixed effect of SOA, allowing to test for both linear and quadratic trends simultaneously. This analysis revealed a significant linear trend (*b* = -20.22, SE = 2.30, *z* = -8.81, *p* < .001) and a non-significant quadratic trend (*b* = 1.81, SE = 2.25, *z* = .80, *p* = .421), indicating that target visibility decayed monotonically as the interval between the tone and target increased. The negative beta coefficient for the linear trend confirms that as the SOA increased (i.e., as the tone was presented later relative to the target), the proportion of seen targets decreased.

#### 3.1.2 Signal Detection Theory measures

The ANOVA for perceptual sensitivity (*d*^′^) demonstrated a significant main effect of SOA (*F*(5, 175) = 9.803, *p* < .001, 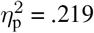). The post-hoc pairwise comparisons (Table S4) revealed higher sensitivity if the alerting tone was presented before the Gabor stimulus was displayed (Figure 3). Further analyses confirmed that *d*^′^ values could be explained by a significant linear trend (*b* = -1.946, SE = .291, *t* = -6.680, *p* < .001), with sensitivity progressively declining as the interval between the tone and target increased. The quadratic trend was not significant (*b* = .118, SE = .291, *t* = .406, *p* = .686).

**Figure 3.**
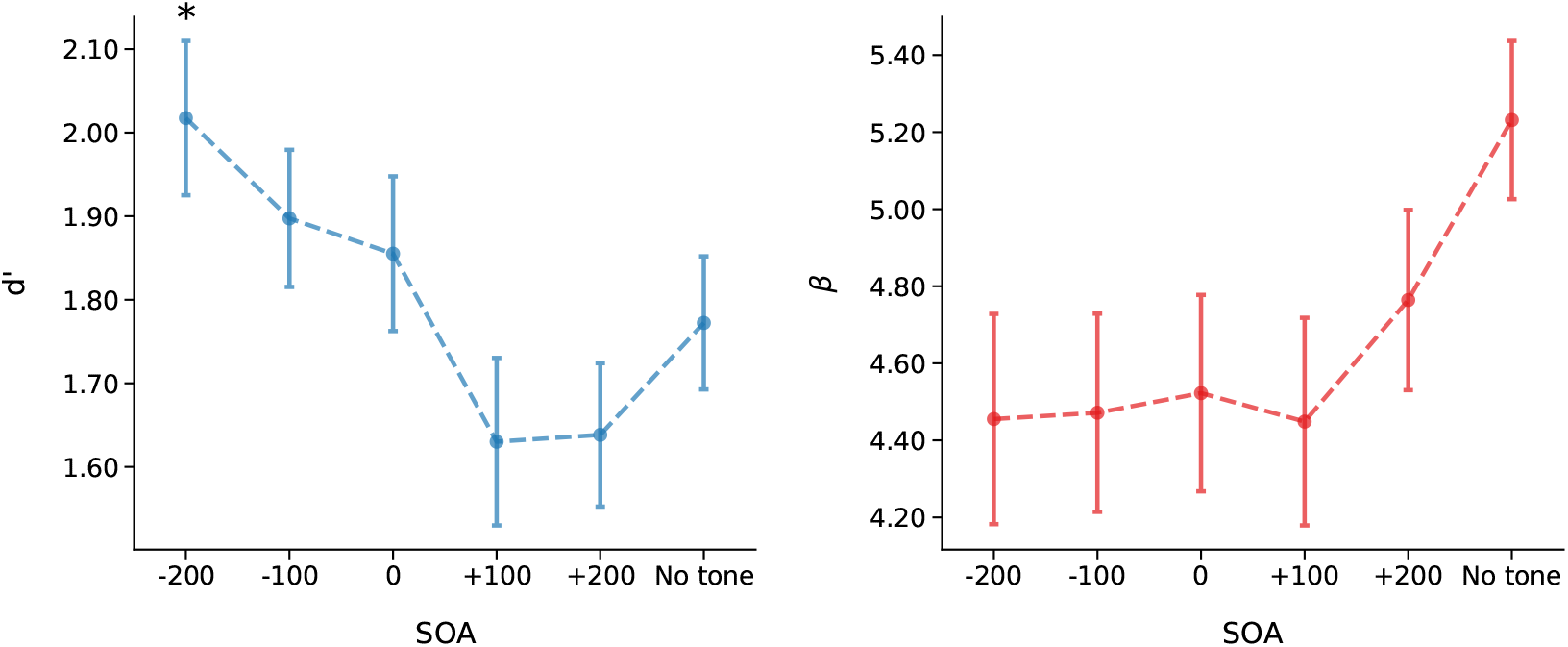
Perceptual sensitivity (*d*^′^) and response criterion (*β*) for Experiment 1 as a function of SOA. Error bars represent standard errors. Asterisks indicate significant post-hoc comparisons against the No tone condition.

**Figure 4.**
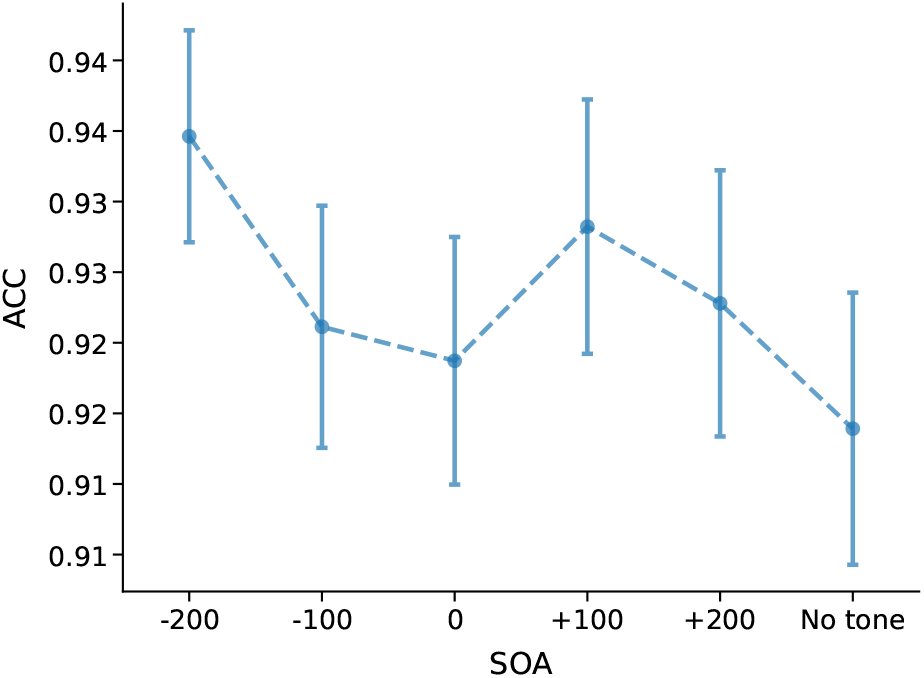
Accuracy for Experiment 1 as a function of SOA. Error bars represent standard errors.

For response criterion (*β*), the main effect of SOA was not significant (*χ*^2^(5) = 3.414, *W* = .019, *p* > .05). Response criterion values did not show significant linear (*b* = 1.129, SE = 1.330, *t* = .849, *p* > .05) or quadratic trends (*b* = -.137, SE = 1.330, *t* = .571, *p* > .05).

#### 3.1.3 ACC

As demonstrated by the t-tests, the proportion of seen targets was higher when the response to the tilt discrimination response was correct, as compared to incorrect responses (*t*(35) = 18.053, *p* < .001, Cohen’s *d* = 2.894, BF_10_ = 1.769 ×10^16^). The LMMs revealed a significant effect of SOA on ACC (*χ*^2^(5) = 11.604, *p* < .05), although pairwise comparisons did not reach significance (see Table S2).

### 3.2 Experiment 2

Experiment 1 confirmed that while pre-cues enhanced performance, no significant retro-cue effects were observed. We hypothesized that the fixed fixation in the previous design acted as a reliable temporal warning signal, allowing participants to precisely predict the Gabor stimulus onset regardless of the alerting cue. To better dissociate general alerting from specific temporal preparation, Experiment 2 introduced temporal uncertainty by using a variable fixation duration ranging between 1000 ms and 1500 ms. This change was implemented to explore how the alerting effect manifests when participants cannot precisely predict the Gabor stimulus onset, and was motivated by findings in the literature suggesting that alerting cues act differently based on stimulus predictability. For fixed-duration stimuli, alerting cues produce effects that are solely associated with increases in the general activation of the system (Fan et al., 2002). However, when stimulus duration varies, these cues exert a stronger influence by providing both a general system activation and critical information regarding the timing of stimulus presentation (Correa et al., 2004; Fan et al., 2002; Kusnir et al., 2011). To test this, Experiment 2 maintained the same design as Experiment 1, but the fixation point had a variable duration between 1000 ms and 1500 ms, rather than a fixed duration. The SOAs used in this experiment were the same as Experiment 1: 200 ms, 100 ms, 0 ms, +100 ms, and +200 ms.

#### 3.2.1 Proportion seen

For the proportion of seen targets, the t-tests showed a significant difference between correct and incorrect trials (*t*(34) = 83.209, *p* < .001, Cohen’s *d* = 1.115, BF_10_ > 1000), with correct trials having a higher proportion of seen responses than incorrect trials. Average FA across participants was .063 (SD = .061).

The LMMs demonstrated a significant effect of SOA (*χ*^2^ = 204.947, *p* < .001). To explore the interaction, post-hoc pairwise comparisons were conducted (see Table S7). Presenting the tone in the pre-stimulus and simultaneous conditions significantly increased the proportion of seen responses compared to the No tone condition. Polynomial contrast analyses revealed significant linear (*b* = -28.008, SE = 2.113, *z* = -13.257, *p* < .001) and quadratic trends (*b* = -9.988, SE = 2.177, *z* = -4.587, *p* < .001).

#### 3.2.2 Signal Detection Theory measures

In the perceptual sensitivity (*d*^′^) analysis, the ANOVA demonstrated a main effect of SOA (*F*(5, 170) = 3.541, *p* < .05,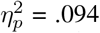), although post-hoc pairwise comparisons did not reach significance (see Table S8). A polynomial contrast analysis confirmed that this effect reflected a linear trend (*b* = -1.100, SE = .364, *p* < .05), with a decrease of the perceptual sensitivity index as the tone was presented after the Gabor (see Figure 6). The model with a quadratic contrast, however, did not significantly fit the data (*b* = -.660, SE = .364, *p* > .05).

**Figure 5.**
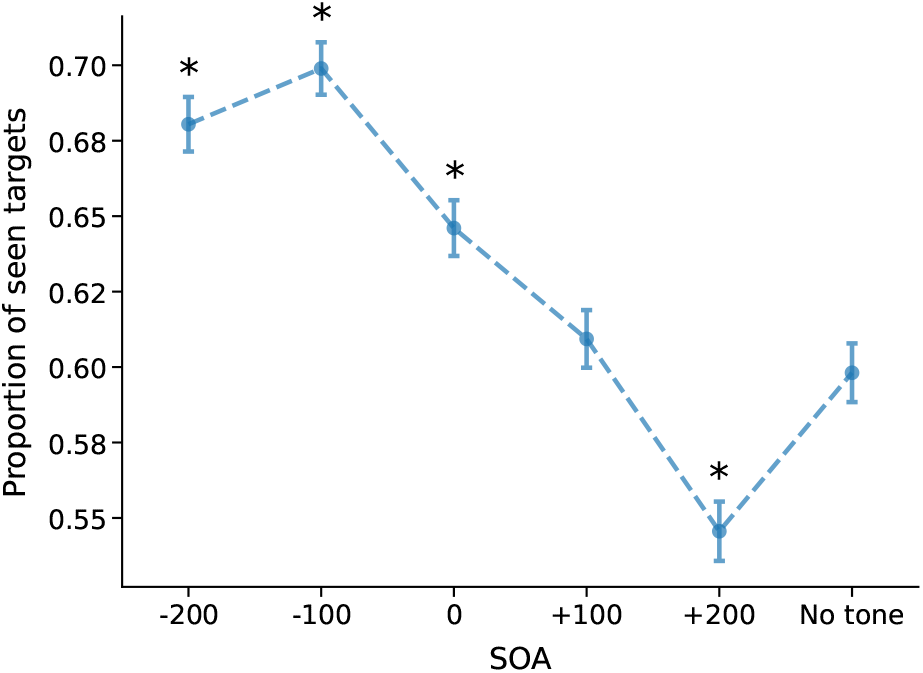
Proportion of seen responses across SOA conditions in Experiment 2. Error bars represent standard errors. Asterisks indicate significant post-hoc comparisons against the No tone condition.

**Figure 6.**
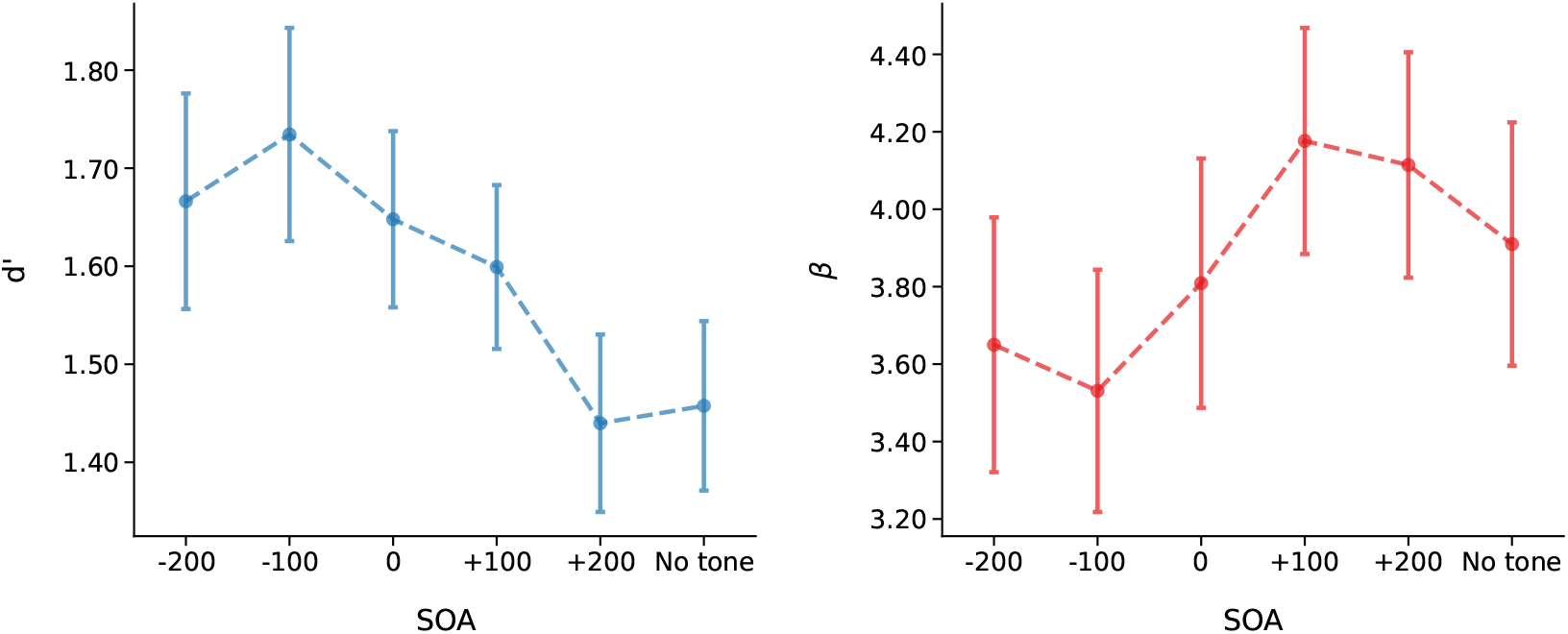
Perceptual sensitivity (*d*^′^) and response criterion (*β*) for Experiment 2 as a function of SOA. Error bars represent standard errors.

**Figure 7.**
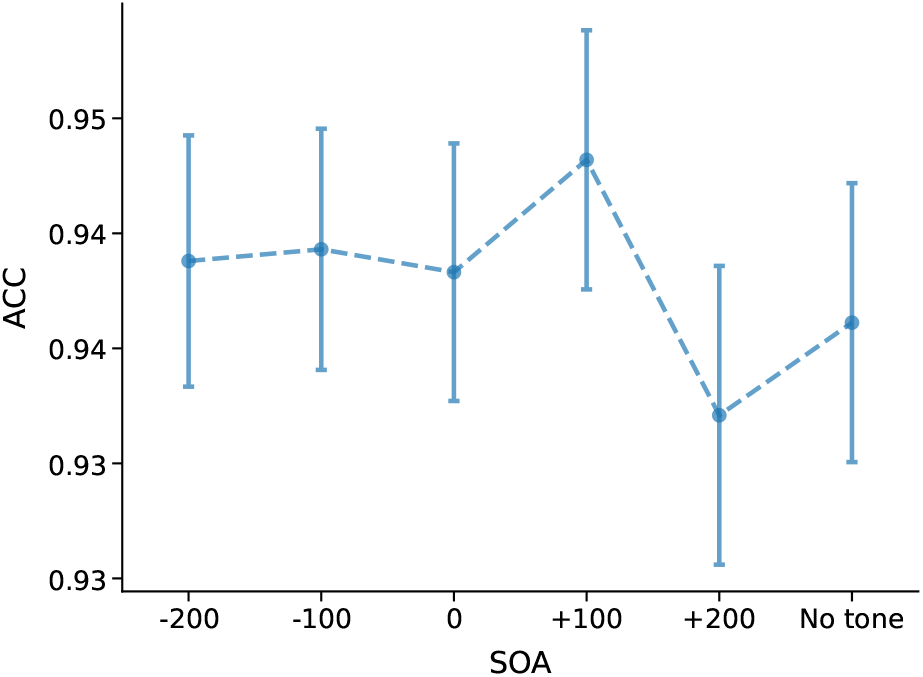
Accuracy across SOA and Awareness conditions in Experiment 2. Error bars represent standard errors.

For the response criterion (*β*), the main effect of SOA was significant (*χ*^2^(5) = 13.171, *W* = .075, *p* < .05), although no post-hoc comparison reached significance (all *p*s > .05; see Table S9). The polynomial contrast analysis revealed a significant linear trend (*b* = 2.945, SE = 1.341, *p* < .05), indicating a more conservative response criterion as the tone was presented later in time (see Figure 6). The quadratic contrast was not significant (*b* = .321, SE = 1.341, *p* > .05).

#### 3.2.3 ACC

Paired-samples t-tests revealed that the seen trials had significantly higher ACC than unseen trials (*t*(34) = 16.427, *p* < .001, Cohen’s *d* = 3.418, BF_10_ = 5.318 × 10^14^). The LMMs analysis of ACC revealed that the SOA was not a significant predictor (*χ*^2^(5) = 5.728, *p* = .333; see Figure S4).

### 3.3 Experiment 3

Although Experiment 2 introduced temporal uncertainty, it did not yet yield significant retro-cue effects. To further investigate the potential for these effects and test the generalizability of the findings across a wider range of time intervals, the variable duration of the fixation cross was maintained in this following experiment, but the SOAs were increased to include a wider range of time intervals. Instead of presenting the tone at ±100 ms and ±200 ms, here the tone could be presented at ±400 ms and ±200 ms, as well as simultaneously (0 ms) or not at all (No tone). This change was motivated by the lack of retro-cue effects even with short SOAs (100 ms) and aimed to explore the effects of alerting tones on visual attention across a broader temporal range, making the design more similar to previous studies (Rimsky-Robert et al., 2019; Sergent, 2018; Sergent et al., 2011, 2013; Thibault et al., 2016). These previous studies also employed larger SOAs ranges, which has been shown to determine the attentional effects that could be found in the task (Milliken et al., 2003).

#### 3.3.1 Proportion seen

The paired-samples t-test revealed a higher proportion of seen targets in correct trials compared to incorrect trials (*t*(34) = 21.902, *p* < .001, Cohen’s *d* = 4.078, BF_10_ = 3.00 *×* 10^18^). The mean proportion of FA was .076 (SD = .122).

In the LMMs analysis a significant main effect of SOA was found (*χ*^2^ (5) = 184.740, *p* < .001). As shown in Figure 8, the proportion of seen responses was significantly higher when the tone was presented before or simultaneously with target onset, compared to when it was presented after the target or when no tone appeared. Post-hoc comparisons (Table S12) revealed that presenting the tone after target onset did not significantly differ from the No tone condition.

**Figure 8.**
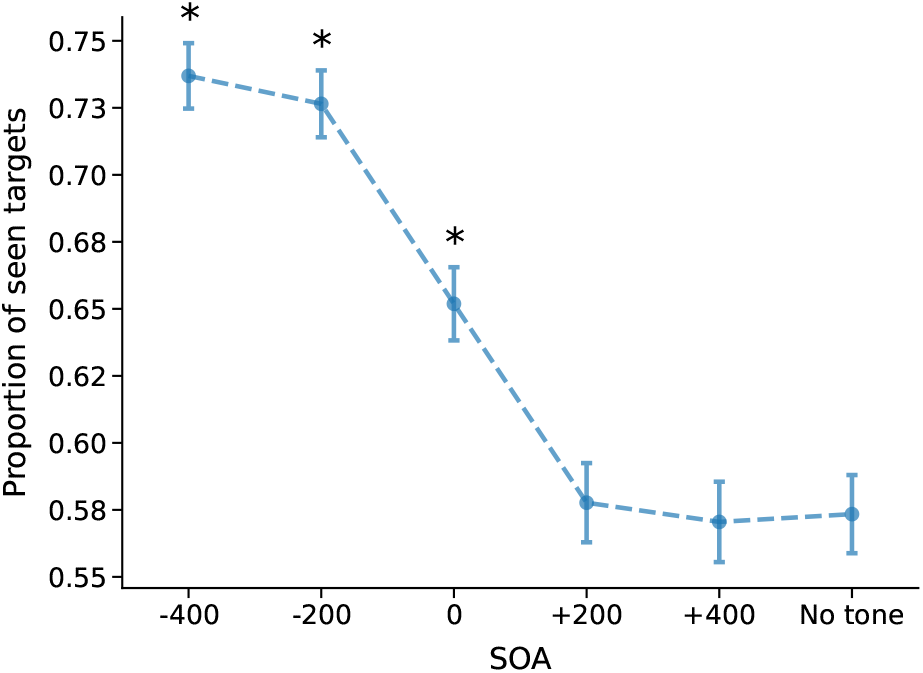
Proportion of seen targets across SOA conditions in Experiment 3. Error bars represent standard errors. Asterisks indicate significant post-hoc comparisons against the No tone condition.

To further explore the pattern of results, polynomial contrasts were conducted. These revealed a significant linear trend (*b* = -26.356, SE = 2.194, *z* = -12.015, *p* < .001), while the quadratic trend was not significant (*b* = .401, SE = 2.206, *z* = .182, *p* = .856).

#### 3.3.2 Signal Detection Theory measures

The analysis of *d*^′^ showed a significant effect of SOA (*χ*^2^(5) = 64.566, *W* = .369, *p* < .001). Post-hoc comparisons (Table S13) revealed that *d*^′^ was significantly higher in all pre-stimulus and simultaneous conditions compared to the post-stimulus and No tone conditions. When the tone was presented after the target onset, there were no differences in perceptual sensitivity when compared to not presenting the tone at all. A polynomial contrast analysis confirmed that this effect reflected a linear trend (*b* = -2.123, SE = .329, *p* < .001), with sensitivity progressively declining as the interval between the tone and target increased (see Figure 9). The model with a quadratic contrast, however, did not significantly fit the data (*b* = .193, SE = .329, *p* = .559).

**Figure 9.**
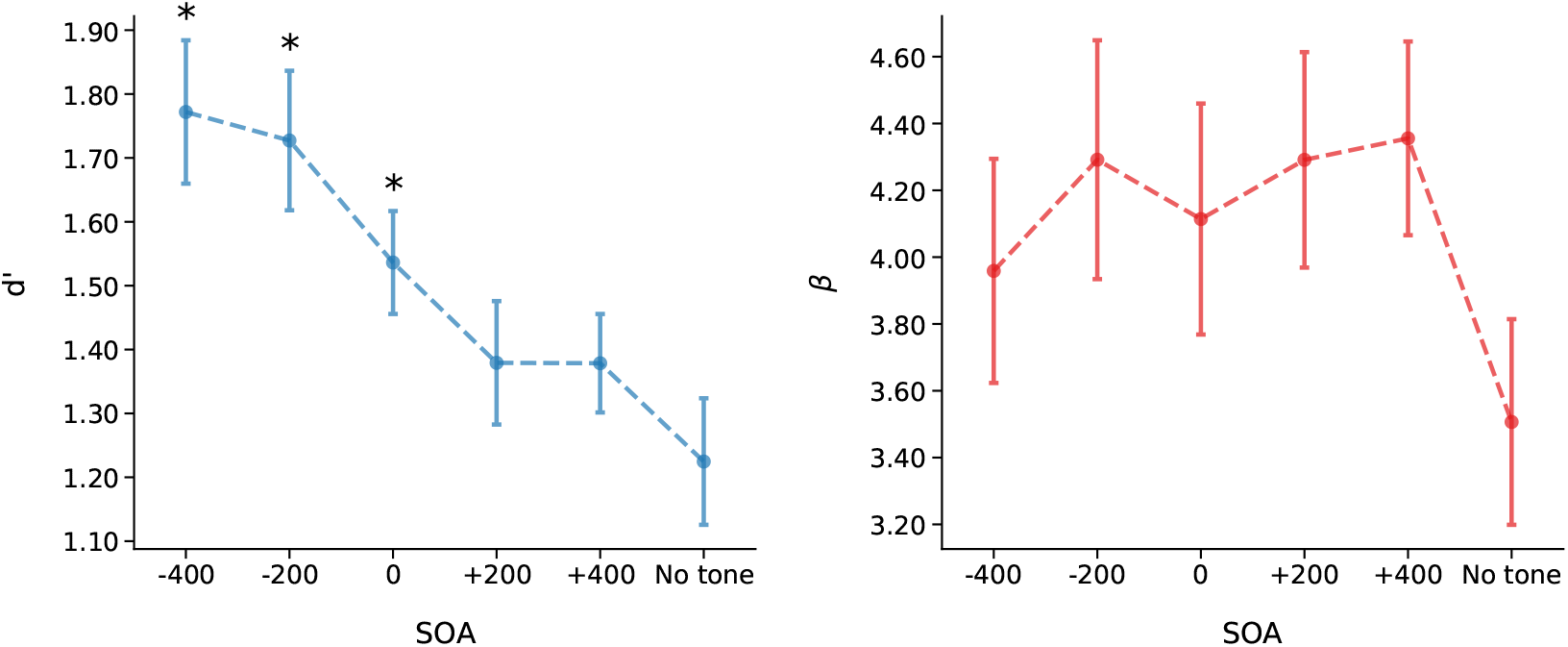
Perceptual sensitivity (*d*^′^) and response criterion (*β*) across SOA conditions in Experiment 3. Error bars represent standard errors. Asterisks indicate significant post-hoc comparisons against the No tone condition.

**Figure 10.**
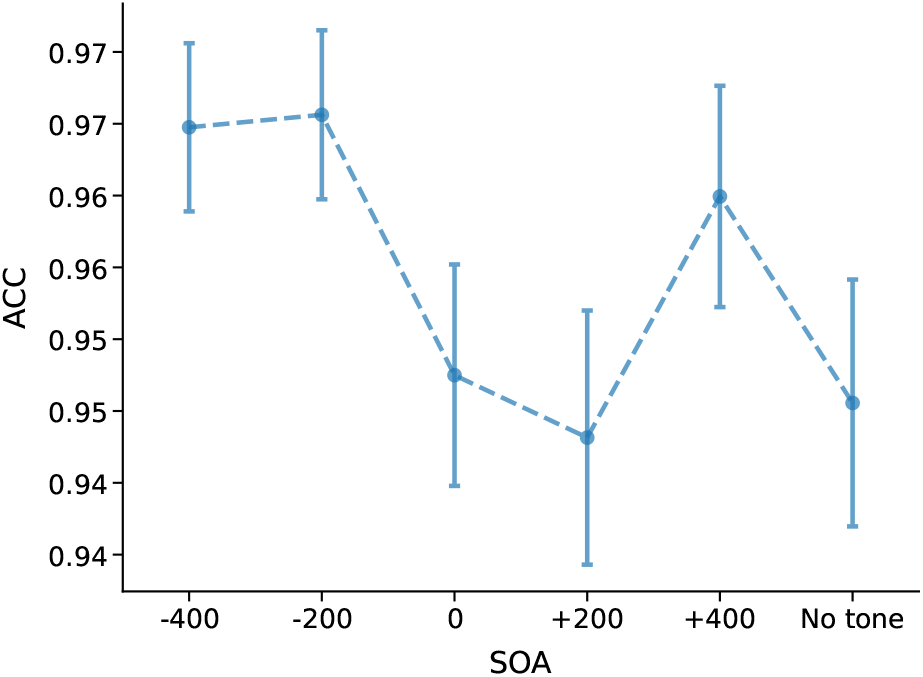
ACC across SOA conditions in Experiment 3. Error bars represent standard errors.

In regards to the response criterion (*β*), the ANOVA revealed a significant effect of SOA (*χ*^2^(5) = 11.367, *W* = .065, *p* < .05), with all SOA conditions showing higher *β* values than the No tone condition, although the post-hoc comparisons did not yield any significant differences (all *p*s > .05; see Table S14). Polynomial contrast analyses showed that neither a linear trend (*b* = 1.484, SE = 1.215, *p* = .224) nor a quadratic trend (*b* = -.288, SE = 1.215, *p* = .813) significantly fit the data.

#### 3.3.3 ACC

The paired-samples t-test showed significant differences in ACC across Awareness levels, with higher accuracy in the seen trials compared to unseen trials (*t*(34) = 30.678, *p* < .001, Cohen’s *d* = 6.089, BF_10_ = 1.16 × 10^23^). In the LMM analysis, the backwards LRTs found no significant effect of SOA (*χ*^2^ (5) = 8.191, *p* > .05).

### 3.4 Experiment 4

Experiment 4 maintained the variable fixation duration (1000 ms - 1500 ms) and the same SOAs as Experiment 3 (−400 ms, −200 ms, 0 ms, +200 ms, and +400 ms), but aimed to investigate whether the effect of retro-cues is modulated by target visibility. This change was motivated by the lack of significant retro-cue effects in the previous experiments, which suggested that at a 50% threshold, the sensory evidence was too weak to be sustained or influenced by retro-cues. According to the Global Neuronal Workspace Theory (GNWT), a sufficient level of bottom-up saliency is a functional requirement for a stimulus to achieve conscious access and trigger global broadcast (Dehaene, 2001; Dehaene and Changeux, 2011; Dehaene et al., 2006; Mashour et al., 2020). We hypothesized that by increasing the visibility to 75%, the higher bottom-up strength would result in a more robust sensory representation. This increased saliency makes it more likely for the system to recover the sensory trace of the stimulus, allowing an auditory retro-cue to provide the necessary amplification to push the fading representation into the global workspace even after the stimulus has disappeared.

#### 3.4.1 Proportion seen

The paired-samples t-test revealed a higher proportion of seen targets in correct trials compared to incorrect trials (*t*(34) = 28.200, *p* < .001, Cohen’s *d* = 6.812, BF_10_ = 7.994 *×* 10^21^). The mean FA rate was .053 (SD = .067).

For the LMM, backwards LRTs revealed a significant main effect of SOA (*χ*^2^ (5) = 197.719, *p* < .001). Post-hoc pairwise comparisons (see Figure 11 and Table S17) showed the highest proportion of seen trials in the pre-stimulus conditions, followed by the simultaneous condition. In the post-stimulus conditions, the +200 ms SOA showed a significantly higher proportion of seen trials compared to the No tone condition, while no significant differences were observed when compared to the +400 ms SOA condition. This late SOA did not differ significantly from the No tone condition either. , the +200 ms effect survived Bonferroni correction for multiple comparisons.

**Figure 11.**
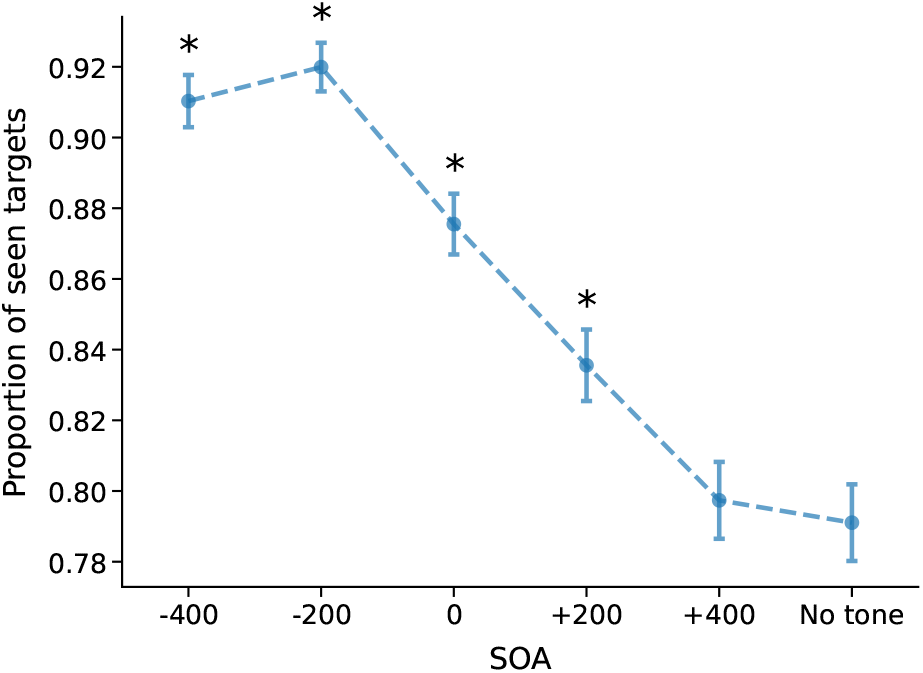
Proportion of Seen responses across SOA conditions in Experiment 4. Error bars represent standard errors. Asterisks indicate significant post-hoc comparisons against the No tone condition

To check for linearity, a polynomial contrast analysis was conducted, revealing a significant linear trend (*b* = -33.418, SE = 3.234, *p* < .001) and a non-significant quadratic component (*b* = -4.934, SE = 3.299, *p* = .135). This further illustrates a decreasing trend in the proportion of seen trials as the tone is presented later in time relative to the target stimulus.

To more directly examine whether the retro-cue effect depended on target visibility, we additionally compared Experiments 3 and 4 in a combined mixed-effects model with SOA and Experiment as fixed factors. This analysis revealed a significant main effect of Experiment, with higher overall proportions of seen reports in Experiment 4 (.857) than in Experiment 3 (.644), and a small but significant SOA * Experiment interaction (*χ*^2^(5) = 11.36, *p* = .045), indicating that the pattern of seen responses across SOAs differed between the 50% and 75% visibility conditions. However, when isolating the retro-cue benefit (proportion seen at each SOA minus the No tone condition), the SOA * Experiment interaction did not reach significance (*p* = .060), although a planned comparison at +200 ms showed a numerically larger benefit in Experiment 4 than in Experiment 3 that again fell short of conventional significance (*p* = .235). These findings therefore provide trend-level, rather than definitive, support for a visibility-dependent modulation of the retro-cue effects. Full details of this between-experiment analysis are reported in the Supplementary Materials.

#### 3.4.2 Signal Detection Theory measures

For *d*^′^, the repeated-measures ANOVA revealed a significant main effect of SOA (F(5, 170) = 8.146, p < .001,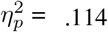). Pairwise comparisons indicated that *d*^′^ was significantly higher when the tone was presented at pre-stimulus and simultaneous SOAs compared to the post-stimulus SOAs and the No tone condition (see Figure 12 and Table S18). This was further confirmed by a polynomial contrast analysis, which revealed a significant linear trend (*b* = -2.309, SE = .542, *p* < .001) and a non-significant quadratic trend (*b* = -0.436, SE = .542, *p* = .422), indicating that sensitivity decreased linearly as the tone was presented in the later SOAs.

**Figure 12.**
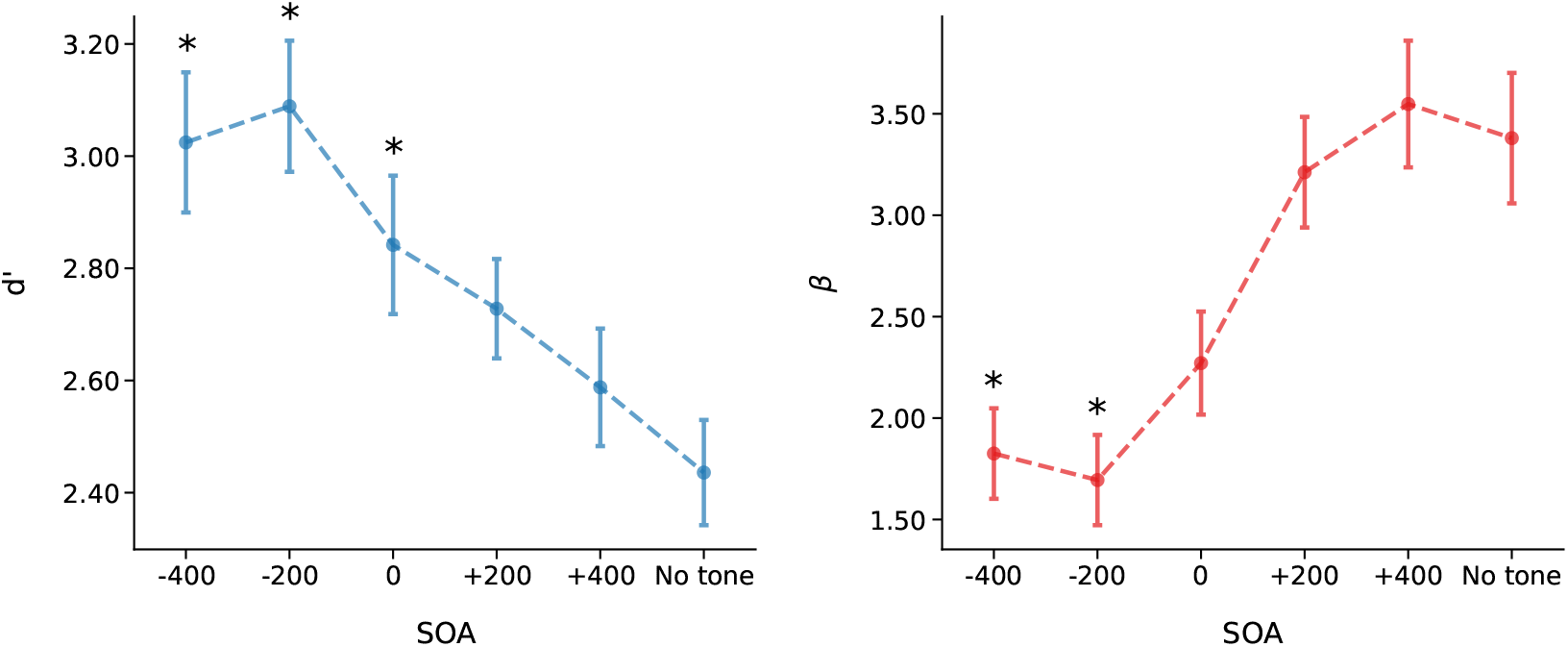
Perceptual sensitivity (*d*^′^) and response criterion (*β*) across SOA conditions in Experiment 4. Error bars represent standard errors. Asterisks indicate significant post-hoc comparisons against the No tone condition

**Figure 13.**
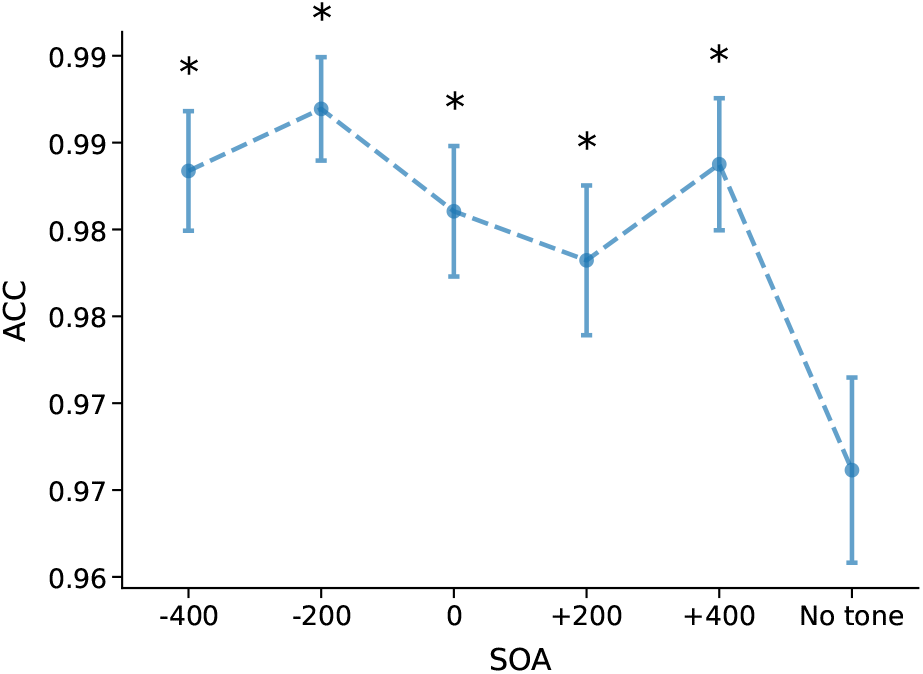
ACC across SOA conditions in Experiment 4. Error bars represent standard errors. Asterisks indicate significant post-hoc comparisons against the No tone condition.

For *β*, the Friedman test revealed a significant main effect of SOA (*χ*^2^(5) = 53.876, *W* = .308, *p* < .001). Post-hoc pairwise comparisons (Table S19) indicated that response criterion became more conservative as the tone was presented later in time relative to the target stimulus (Figure 12). Polynomial contrast analysis for *β* showed a significant linear trend (*b* = 9.286, SE = 1.165, *p* < .001) and a non-significant quadratic trend (*b* = 2.052, SE = 1.165, *p* = .080), indicating a linear increase in response criterion as the tone was presented later in time relative to the target stimulus.

#### 3.4.3 ACC

Paired-samples t-tests comparing accuracy between seen and unseen trials revealed significantly higher accuracy for seen trials compared to unseen trials (*t*(34) = 23.786, *p* < .001, Cohen’s *d* = 5.130, BF_10_ = 3.852 × 10^19^). For the LMM analysis, backwards LRTs revealed a significant main effect of SOA (*χ*^2^ (5) = 12.220, *p* = .031). Post-hoc pairwise comparisons indicated a general effect of the tone increasing the ACC when compared to no presenting the tone, regardless of the SOA (see Figure S8 and Table S16).

## 4 Discussion

The present set of experiments investigated whether and how the temporal dynamics of an alerting tone, presented at various SOAs, modulates conscious perception of a centrally presented Gabor stimulus. Based on evidence that attention often serves as a gateway to consciousness, this research focused on the alerting network, which maintains vigilance and responds to environmental changes. While phasic alerting (transient increase in alertness) has been shown to enhance conscious perception when cues precede the target (Botta et al., 2014; Chica et al., 2016; Kusnir et al., 2011; A. Petersen et al., 2017; Poth, 2025), the effects of auditory retro-cues have been less clear. The studies dealing with retro-cues usually involve visual cueing (Gressmann and Janczyk, 2016; Sergent, 2018; Szaszkó et al., 2024; Thibault et al., 2016; Xia et al., 2016; Zerr et al., 2021), and even when employing auditory cues they usually involve lateralization of either the stimulus, introducing spatial uncertainty, or the cues, which could also be lateralized (Gressmann and Janczyk, 2016; Janczyk and Reuss, 2016; Rimsky-Robert et al., 2019; Sergent, 2018; Sergent et al., 2013; Szaszkó et al., 2024; Thibault et al., 2016; Xia et al., 2016). These experimental designs introduce confounding factors, making it difficult to isolate the specific effects of alerting and spatial attention on conscious perception.

Across four experiments, we systematically manipulated the timing of an auditory alerting cue relative to the onset of a near-threshold Gabor stimulus, and assessed its effects on perceptual awareness and SDT indices. All four experiments employed a visual detection task where participants reported the presence or absence of a low-contrast Gabor patch, paired with an auditory alerting cue presented at different time intervals (before, simultaneously, or after relative to Gabor onset). Each experiment also included a no-tone control condition.

Our findings consistently revealed that, when presented before target onset, alerting cues provided robust benefits for both subjective awareness (increased proportion of seen targets) and objective perceptual sensitivity (higher d’) compared to the no-tone control. These results support existing literature indicating that pre-stimulus phasic alerting enhances the conscious perception of near-threshold stimuli by transiently boosting neural excitability or attentional resources at critical moments (Fan et al., 2005; Sadaghiani et al., 2010; Thiel and Fink, 2007). These effects have been found to involve increased visual processing speed, lowered thresholds for conscious perception, and the efficient allocation of attentional resources, often reflected in physiological markers such as pupil diameter (Feng et al., 2017; Kusnir et al., 2011; A. Petersen et al., 2017).

A primary goal of this study was to explore the effects of auditory retro-cues. Contrary to the robust enhancing effects of pre-stimulus cues, auditory alerting tones presented after target onset did not significantly enhance conscious perception (proportion of seen targets or d’) compared to the No tone condition in Experiments 1, 2, and 3. While some studies suggest retro-cues can enhance perception of past stimuli (Garnier-Allain et al., 2023), the evidence is richer for visual cues (Gressmann and Janczyk, 2016; Sergent, 2018; Szaszkó et al., 2024; Thibault et al., 2016; Xia et al., 2016; Zerr et al., 2021). One possible explanation for the lack of significant retro-cue effects in the first three experiments is the low visibility threshold employed (∼50%). According to the GNWT, conscious access is a non-linear process that requires a stimulus to possess sufficient bottom-up saliency to trigger “global ignition”, a large-scale broadcast of information across a fronto-parietal network (Dehaene and Changeux, 2011; Dehaene et al., 2006). In Experiments 13, the near-threshold contrast likely generated sensory representations that lacked the necessary strength to be sustained. Within the GNWT framework, such weak signals decay rapidly, failing to reach the stability required for successful late amplification by post-stimulus signals (Mashour et al., 2020). Consequently, by the time the retrospective alerting tone was presented, the sensory trace of the Gabor stimulus may have already dissipated, making it unavailable for the non-linear amplification required for awareness. By increasing visibility to 75% in Experiment 4, we likely provided a more robust bottom-up signal that remained available for later modulation, aligning with findings from successful retro-perception studies that utilize higher baseline accuracies (Rimsky-Robert et al., 2019).

Other factors likely contributed to the absence of effects in the initial experiments. The high cognitive demand of performing two tasks (detection and discrimination) under near-threshold conditions may have led to a misallocation of attentional resources (Causse et al., 2022; Mandal et al., 2024; Poth, 2020; Weinbach and Henik, 2012b). Under such taxing conditions, retro-cues can even introduce dual-task costs that impair working memory processes (Katus and Eimer, 2019; Souza and Oberauer, 2017; Zickerick et al., 2020), and in turn compromise conscious access. Additionally, the influence of backward masking cannot be ignored; a later stimulus can disrupt the processing of a weak target through cortical suppression (Mattingly et al., 2018; Silverstein, 2015). In our design, these interference effects may have been particularly strong at the 50% threshold, preventing the retro-cue from exerting a beneficial influence.

Experiment 4 demonstrated that the retro-cue effect was observable under higher-visibility conditions; by increasing the stimulus contrast to achieve a 75% detection rate, we found that the +200 ms SOA condition significantly increased the proportion of seen reports compared to the No tone condition. This suggests that, under higher visibility conditions, non-spatial auditory retro-cues can modulate the likelihood of seeing a target, although the cross-visibility comparison between Experiments 3 and 4 only provided trend-level, rather than definitive, evidence for a visibility-dependent modulation of this effect. It is important to note, however, that in Experiment 4 the target was already seen on the majority of trials, meaning that some targets in the +200 ms condition may have already been partially available before the tone arrived. We therefore cannot rule out that the observed increase in seen reports reflects a boost in confidence or reportability rather than a transition of an unconscious representation into awareness. Disentangling these possibilities would require paradigms that more precisely characterise the state of the sensory representation at the time of cue onset.

In the present data set, the SDT analyses were not sensitive enough to determine whether the +200 ms benefit reflects a genuine increase in sensory evidence or a shift in decision criterion. The SOA-dependent variation in response criterion in Experiment 4 makes a purely perceptual interpretation less straightforward. Our data showed that response criterion increased linearly as the tone occurred later in time, indicating that participants adopted a progressively more conservative reporting strategy in the retro-cue conditions. This means that the increase in seen reports at +200 ms emerged despite a more conservative criterion at that SOA relative to the pre-stimulus conditions, which in principle argues against a simple liberal-bias account of the retro-cue benefit. However, because response criterion did not differ significantly between the +200 ms and No tone conditions in the post-hoc comparisons, we cannot fully rule out that the increase in seen reports at +200 ms partially reflects criterion changes rather than purely perceptual changes. Future studies could combine pre- and retro-cue paradigms with graded awareness or confidence scales (e.g., the Perceptual Awareness Scale or trial-wise confidence ratings) and type-2 SDT or meta-cognitive modelling, to more directly test whether alerting retro-cues primarily modulate preconscious sensory traces, phenomenally conscious but inaccessible representations, or post-perceptual decision processes.

A further methodological consideration concerns the order of responses. Across all experiments, participants first made the orientation discrimination response and only afterwards reported whether they had seen the Gabor or not. This order was chosen to prevent the visibility judgment from biasing the discrimination response, following standard practice in near-threshold detection paradigms (Chica et al., 2016; Rodríguez-San Esteban et al., 2025; Sergent et al., 2013; Thibault et al., 2016). However, it introduces the possibility that confidence in the discrimination response could have influenced the subsequent seen/unseen judgment: participants who felt they had successfully discriminated the orientation may have been more likely to report the stimulus as seen. Indeed, dissociations between discrimination confidence and subjective visibility have been reported in near-threshold tasks (Rausch and Zehetleitner, 2016), suggesting that the two measures are not fully interchangeable. We cannot rule out this response-order effect, and it should be borne in mind when interpreting the seen reports as a direct index of conscious access. Notably, if such an influence were present, it would apply equally across all SOA conditions and would therefore not account for the SOA-specific effects observed, particularly the increase in seen reports at +200 ms in Experiment 4. Future work could counterbalance or reverse the response order to evaluate the robustness of the seen/unseen measure independently of the discrimination response.

Additionally, the effects of retro-cues in Experiment 4 extended beyond subjective awareness to objective discrimination performance. While Experiments 1 through 3 showed limited or non-significant impacts on orientation discrimination accuracy, Experiment 4 demonstrated that every SOA condition significantly improved accuracy compared to the no-tone control. This generalized benefit across all timing conditions, including both pre and retro-cues, suggests that once target visibility was higher, the auditory tone enhanced discrimination performance regardless of its temporal relationship to the stimulus. The fact that this improvement extended to retro-cue conditions (+200 and +400 ms) is consistent with the interpretation that the system can leverage late alerting signals when the sensory representation is sufficiently strong, though the specific mechanism (whether perceptual enhancement, response bias, or a combination) remains to be determined.

The finding that post-stimulus alerting tones can modulate visibility reports under higher-visibility conditions provides an opportunity to reflect on competing theoretical frameworks of conscious access. Specifically, our results can be interpreted through two distinct lenses: the phenomenal/access distinction proposed by Block (1995, 2005) and the GNWT defended by authors such as Baars (1988) and Dehaene (2001, 2006). As argued by Sergent (2018), the “desynchronization” of conscious access from external stimulation suggests that the timing of awareness is not strictly time-locked to sensory entry, but can be influenced “offline” when a representation becomes relevant for a task. This decoupling of stimulus onset from the timing of conscious access is, as noted by Garnier-Allain et al.(2023), a key point of distinction between current theories of conscious perception, as they offer fundamentally different accounts of the state of the unseen trace prior to the arrival of the cue. Despite these theoretical considerations, our paradigm cannot adjudicate between them: we measured whether the cue changed the proportion of seen reports, not the underlying state of the sensory trace. Further studies employing neuroimaging techniques such as electroencephalography in similar pre- and retro-cueing paradigms could track the time course of stimulus-evoked activity and late cue-related responses, and thereby begin to disentangle preconscious sensory traces, phenomenally conscious but inaccessible representations, and post-perceptual decision-level changes.

In the context of the GNWT, the delay between the stimulus and the cue would be occupied by a preconscious state (Dehaene et al., 2006; Kouider and Dehaene, 2007), a period where the sensory trace is strong enough to be maintained in local recurrent loops within the visual cortex but lacks the necessary top-down amplification to trigger the non-linear “ignition” of the global workspace to reach awareness (Dehaene et al., 2006; Gaillard et al., 2009; Mashour et al., 2020). Within this framework, consciousness is identified with global access; therefore, information that is not broadcasted remains strictly unconscious. Under this account, the auditory cue at +200 ms could act as a late attentional booster that pushes a decaying preconscious trace over the threshold of global broadcasting, which typically involves long-range coordination between fronto-parietal and sensory networks (Dehaene, 2001; Del Cul et al., 2009; Sergent et al., 2013). Conversely, the phenomenal/access distinction posits that the stimulus may have already entered a state of phenomenal consciousness, that is, the rich, qualitative “what it is like” aspect of experience (P) which “overflows” the limited capacity of the access system (A) (Amir et al., 2023; Block, 2005, 2011). The capacity of the phenomenal system is proposed to be significantly larger than that of the working memory buffer required for conscious report (Block, 2011, 2014). In this view, the cue does not “create” the experience post-hoc but rather facilitates access to an existing phenomenal representation by providing the attentional prioritization required for report and the rational control of action (Amir et al., 2023; Block, 2005, 2007; Overgaard, 2018).

The distinction between these models hinges on whether the unseen trace is strictly unconscious (preconscious) or phenomenally conscious but inaccessible. Proponents of the GNWT argue that conscious access corresponds to a non-linear ignition of a distributed fronto-parietal networkincluding the prefrontal cortex, parietal regions, and the anterior cingulatewhich allows information to be globally broadcasted and made available for report (Dehaene, 2001; Dehaene et al., 2006; Mashour et al., 2020). Within this framework, prefrontal activity is considered a critical requirement for an experience to be conscious. In contrast, authors like Ned Block associate phenomenal (P) consciousness with posterior cortical “hot zones” encompassing occipital and temporal sensory areas ((Block, 2005)). From this perspective, these posterior hot zones are sufficient for basic conscious experience, while the fronto-parietal global workspace is specifically tied to access (A) consciousness and the cognitive processes required for reporting (Koch et al., 2016; Nani et al., 2019). Our data show that a non-spatial auditory alert can modulate seen reports for stimuli that have already disappeared, which is consistent with the idea that conscious access is a flexible process not strictly determined at stimulus onset (Del Cul et al., 2009; Sergent, 2018). Whether this reflects a genuine late transition into awareness or a post-perceptual boost in reportability cannot be determined from the present data alone, and both frameworks remain plausible interpretations.

## 5 Conclusions

In summary, these four experiments demonstrate that the timing of an alerting cue critically determines its ability to modulate conscious perception and behavioral performance. Phasic alerting, when optimally timed, increases the likelihood of detecting faint stimuli, highlighting the importance of the alerting system in conscious access. These findings contribute to our understanding of the interplay between alerting, attention, and awareness, and have direct implications for designing interventions to improve perceptual performance in applied settings, such as brain-computer interfaces or neurofeedback systems.

Our results are consistent with prior studies demonstrating that phasic alerting can improve perceptual performance and awareness (Kusnir et al., 2011; Matthias et al., 2010). The present research extends these findings by providing a fine-grained analysis of SOA effects across different experimental designs and participant samples, revealing that alerting benefits are strongest for pre-stimulus and simultaneous cues. Crucially, while retro-cues in the first three experiments did not reliably modulate the proportion of seen targets or perceptual sensitivity, Experiment 4 showed that a tone presented 200 ms after target offset significantly increased seen reports when target visibility was higher (∼75%). This pattern suggests that the effectiveness of alerting retro-cues may be influenced by target visibility and the robustness of the initial sensory representation, rather than reflecting a general capacity to bring near-threshold stimuli into awareness, although our behavioural and SDT measures do not allow us to fully separate perceptual from decisional contributions to the +200 ms effect. Ultimately, the temporal specificity of these effects (being strongest for pre-stimulus and simultaneous cues, yet present for retrospective cues under specific conditions) aligns with models proposing that alerting modulates the accumulation of evidence required for conscious perception (Matthias et al., 2010; A. Petersen et al., 2017), while also highlighting the need for future work to clarify whether the locus of this modulation is perceptual, post-perceptual, or both.

## Supplementary materials

### Experiment 1

#### Reaction times

T-tests were conducted to compare the RTs between Awareness conditions, showing a significant difference between Seen and Unseen trials (*t*(35) = 10.715, *p* < .001, Cohen’s *d* = 2.063, BF_10_) = 6.153 × 10^9^ and demonstrating that RTs were significantly faster in the Unseen condition.

The hierarchical backwards LRTs revealed a significant effect of SOA (*χ*^2^(5) = 14.332, *p* < .05). Post-hoc analyses (see Table S1) showed a trend of faster RTs when the alerting tone was presented before or simultaneous to the Gabor stimulus (see Figure S1), although posthoc comparisons did not reach significance.

**Figure S1.**
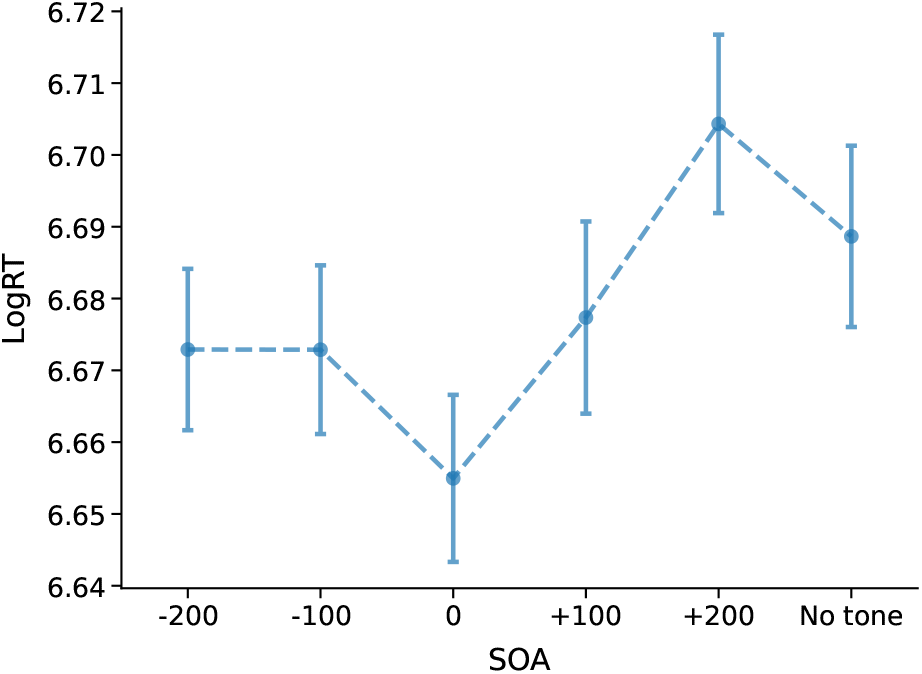
Reaction times for Experiment 1 as a function of SOA. Error bars represent standard errors.

**Table S1.**
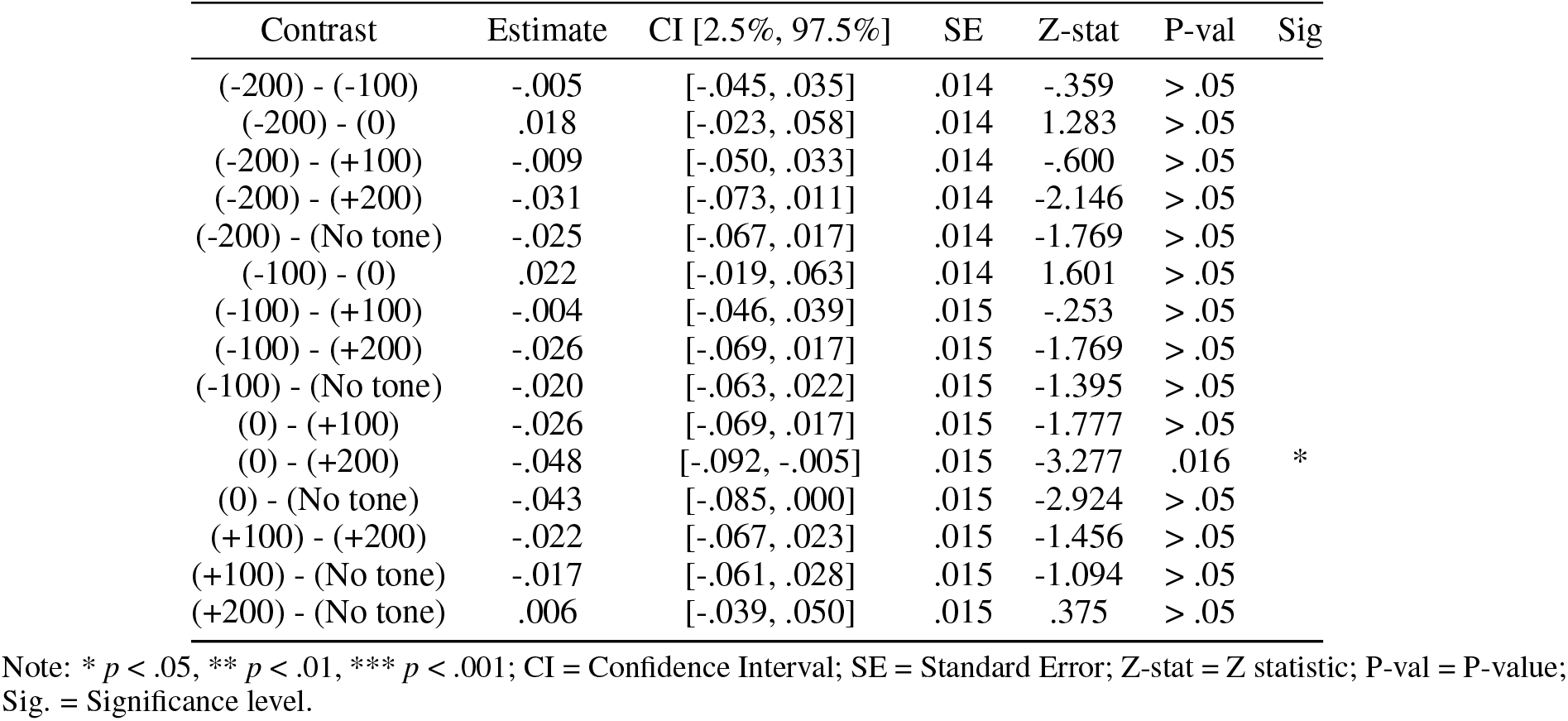
Post-hoc pairwise comparisons of RT for each level of SOA in Experiment 1.

#### Accuracy

As demonstrated by the t-tests, the proportion of seen targets was higher when the response to the tilt discrimination response was correct, as compared to incorrect responses (*t*(35) = 18.053, *p* < .001, Cohen’s *d* = 2.894, BF_10_ = 1.769 *×* 10^16^).

**Figure S2.**
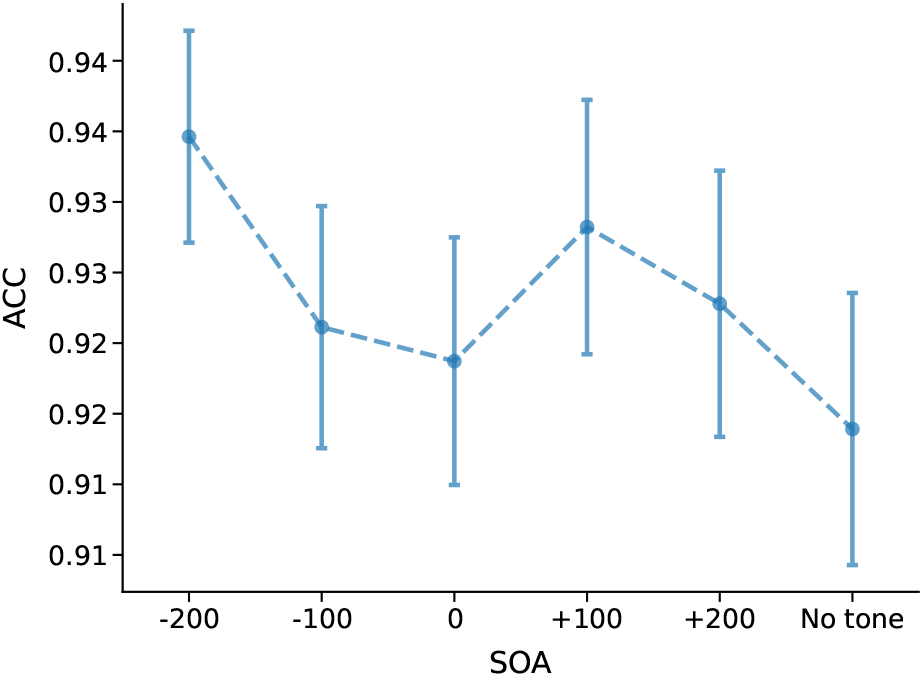
Accuracy for Experiment 1 as a function of SOA. Error bars represent standard errors.

**Table S2.** Post-hoc pairwise comparisons of ACC for each level of SOA in Experiment 1.

| Contrast | Estimate | CI [2.5%, 97.5%] | SE | Z-stat | P-val | Sig |
| --- | --- | --- | --- | --- | --- | --- |
| (-200) - (-100) | .417 | [-.265, 1.099] | .232 | 1.796 | > .05 |  |
| (-200) - (0) | .041 | [-.670, .752] | .242 | .170 | > .05 |  |
| (-200) - (+100) | -.194 | [-.973, .584] | .265 | -.732 | > .05 |  |
| (-200) - (+200) | .263 | [-.463, .990] | .247 | 1.065 | > .05 |  |
| (-200) - (No tone) | .558 | [-.150, 1.265] | .241 | 2.314 | > .05 |  |
| (-100) - (0) | -.376 | [-1.065, .313] | .235 | -1.600 | > .05 |  |
| (-100) - (+100) | -.611 | [-1.372, .149] | .259 | -2.360 | > .05 |  |
| (-100) - (+200) | -.154 | [-.856, .549] | .239 | -.642 | > .05 |  |
| (-100) - (No tone) | .140 | [-.539, .820] | .232 | .606 | > .05 |  |
| (0) - (+100) | -.235 | [-1.016, .545] | .266 | -.885 | > .05 |  |
| (0) - (+200) | .222 | [-.508, .952] | .249 | .893 | > .05 |  |
| (0) - (No tone) | .516 | [-.198, 1.230] | .243 | 2.123 | > .05 |  |
| (+100) - (+200) | .458 | [-.339, 1.254] | .271 | 1.686 | > .05 |  |
| (+100) - (No tone) | .752 | [-.031, 1.534] | .267 | 2.820 | > .05 |  |
| (+200) - (No tone) | .294 | [-.432, 1.020] | .247 | 1.189 | > .05 |  |

Note: \* $p < .05$ , \*\* $p < .01$ , \*\*\* $p < .001$ ; CI = Confidence Interval; SE = Standard Error; Z-stat = Z statistic; P-val = P-value; Sig. = Significance level.

The LMMs revealed a significant effect of SOA on ACC (*χ*^2^(5) = 11.604, *p* < .05), although pairwise comparisons did not reach significance (see Table S2).

### Proportion seen and SDT tables

**Table S3.** Post-hoc pairwise comparisons of Seen for each level of SOA in Experiment 1.

| Contrast | Estimate | CI [2.5%, 97.5%] | SE | Z-stat | P-val | Sig |
| --- | --- | --- | --- | --- | --- | --- |
| (-200) - (-100) | .241 | [-.018, .500] | .088 | 2.727 | > .05 |  |
| (-200) - (0) | .366 | [.110, .623] | .087 | 4.197 | < .001 | *** |
| (-200) - (+100) | .611 | [.352, .870] | .088 | 6.924 | < .001 | *** |
| (-200) - (+200) | .696 | [.438, .955] | .088 | 7.913 | < .001 | *** |
| (-200) - (No tone) | .611 | [.354, .868] | .088 | 6.983 | < .001 | *** |
| (-100) - (0) | .126 | [-.129, .380] | .087 | 1.448 | > .05 |  |
| (-100) - (+100) | .370 | [.113, .627] | .088 | 4.221 | < .001 | *** |
| (-100) - (+200) | .455 | [.199, .712] | .087 | 5.214 | < .001 | *** |
| (-100) - (No tone) | .370 | [.116, .625] | .087 | 4.267 | < .001 | *** |
| (0) - (+100) | .244 | [-.010, .498] | .087 | 2.821 | > .05 |  |
| (0) - (+200) | .330 | [.076, .583] | .086 | 3.820 | .002 | ** |
| (0) - (No tone) | .245 | [-.007, .496] | .086 | 2.851 | > .05 |  |
| (+100) - (+200) | .086 | [-.170, .341] | .087 | .984 | > .05 |  |
| (+100) - (No tone) | .000 | [-.254, .254] | .087 | .004 | > .05 |  |
| (+200) - (No tone) | -.085 | [-.338, .168] | .086 | -.988 | > .05 |  |

Note: \* $p < .05$ , \*\* $p < .01$ , \*\*\* $p < .001$ ; CI = Confidence Interval; SE = Standard Error; Z-stat = Z statistic; P-val = P-value; Sig. = Significance level.

**Table S4.** Post-hoc pairwise comparisons of *d*^′^ for each level of SOA in Experiment 1.

| Contrast | T-stat | DF | P-val | BF10 | Cohen's $d$ | Sig. |
| --- | --- | --- | --- | --- | --- | --- |
| (-100) vs (-200) | -1.834 | 35 | > .05 | .808 | -.229 |  |
| (-100) vs (0) | .631 | 35 | > .05 | .216 | .081 |  |
| (-100) vs (+100) | 3.892 | 35 | .006 | 67.436 | .486 | ** |
| (-100) vs (+200) | 3.825 | 35 | .008 | 56.73 | .514 | ** |
| (-100) vs (No tone) | 1.875 | 35 | > .05 | .863 | .258 |  |
| (-200) vs (0) | 2.696 | 35 | > .05 | 3.991 | .293 |  |
| (-200) vs (+100) | 5.540 | 35 | < .001 | 6009.368 | .670 | *** |
| (-200) vs (+200) | 5.268 | 35 | < .001 | 2797.977 | .709 | *** |
| (-200) vs (No tone) | 3.294 | 35 | .034 | 15.291 | .474 | * |
| (0) vs (+100) | 3.263 | 35 | .037 | 14.2 | .389 | * |
| (0) vs (+200) | 3.125 | 35 | > .05 | 10.293 | .405 |  |
| (0) vs (No tone) | 1.070 | 35 | > .05 | .304 | .160 |  |
| (+100) vs (+200) | -.113 | 35 | > .05 | .18 | -.015 |  |
| (+100) vs (No tone) | -2.205 | 35 | > .05 | 1.52 | -.262 |  |
| (+200) vs (No tone) | -2.105 | 35 | > .05 | 1.269 | -.270 |  |

Note: \* $p < .05$ , \*\* $p < .01$ , \*\*\* $p < .001$ ; CI = Confidence Interval; SE = Standard Error; Z-stat = Z statistic; P-val = P-value; Sig. = Significance level.

### Experiment 2

#### Reaction times

First, paired-samples t-tests were conducted to compare the RTs between the seen and unseen trials. The results showed that the seen trials had significantly lower RTs (*t*(34) = 7.341, *p* < .001, Cohen’s *d* = 1.392, BF_10_ = 8.027 ×10^5^) than the unseen trials.

The hierarchical backwards LRTs revealed a significant effect of SOA (*χ*^2^(5) = 34.547, *p* < .001).

**Figure S3.**
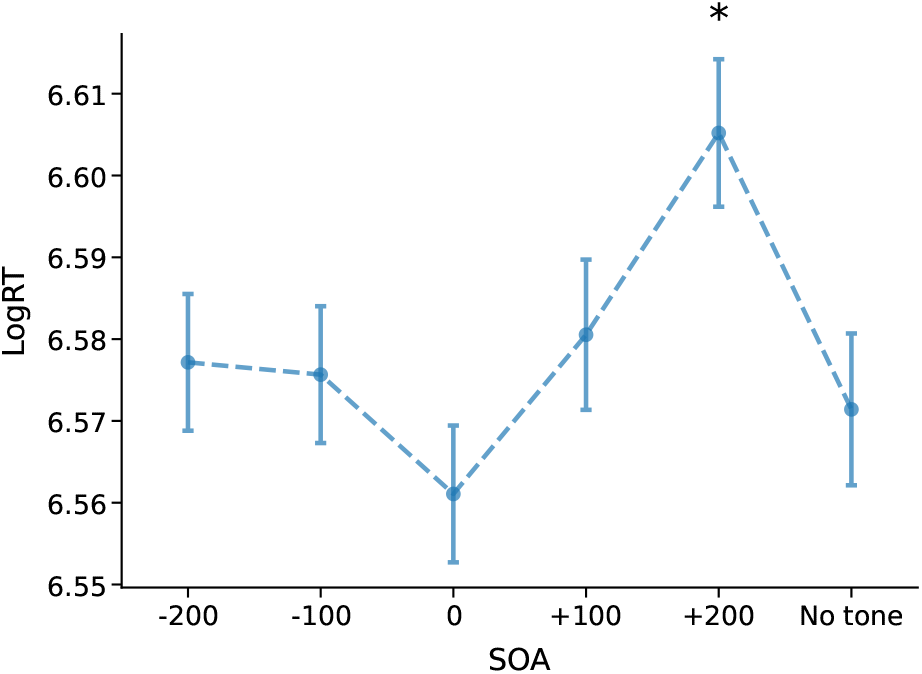
Reaction times (RT) across SOA conditions in Experiment 2. Error bars represent standard errors. Asterisks indicate significant post-hoc comparisons against the No tone condition

Posthoc pairwise comparisons were conducted to further explore this effect (see Table S5). This revealed that although pre-stimulus (-200 and -100 ms), simultaneous (0 ms), and post-stimulus (+100 ms) tones showed significantly faster RTs than the +200 ms condition, they were not significantly different to the no tone condition. The only significant difference was between the +200 ms and No tone conditions, demonstrating that presenting the tone 200 ms after the Gabor stimulus produced significantly slower responses.

**Table S5.** Post-hoc pairwise comparisons of RT for each level of SOA in Experiment 2.

| Contrast | Estimate | CI [2.5%, 97.5%] | SE | Z-stat | P-val | Sig |
| --- | --- | --- | --- | --- | --- | --- |
| (-200) - (-100) | .008 | [-.020, .035] | .009 | .821 | > .05 |  |
| (-200) - (0) | .016 | [-.013, .045] | .010 | 1.651 | > .05 |  |
| (-200) - (+100) | -.008 | [-.037, .022] | .010 | -.756 | > .05 |  |
| (-200) - (+200) | -.041 | [-.071, -.011] | .010 | -3.973 | .001 | ** |
| (-200) - (No tone) | -.005 | [-.035, .024] | .010 | -.510 | > .05 |  |
| (-100) - (0) | .008 | [-.020, .036] | .010 | .871 | > .05 |  |
| (-100) - (+100) | -.015 | [-.044, .013] | .010 | -1.561 | > .05 |  |
| (-100) - (+200) | -.049 | [-.079, -.019] | .010 | -4.786 | < .001 | *** |
| (-100) - (No tone) | -.013 | [-.042, .016] | .010 | -1.299 | > .05 |  |
| (0) - (+100) | -.024 | [-.053, .006] | .010 | -2.343 | > .05 |  |
| (0) - (+200) | -.057 | [-.088, -.026] | .010 | -5.465 | < .001 | *** |
| (0) - (No tone) | -.021 | [-.051, .009] | .010 | -2.079 | > .05 |  |
| (+100) - (+200) | -.034 | [-.065, -.002] | .011 | -3.168 | .023 | * |
| (+100) - (No tone) | .002 | [-.028, .033] | .010 | .229 | > .05 |  |
| (+200) - (No tone) | .036 | [.004, .067] | .011 | 3.334 | .013 | * |

Note: \* $p < .05$ , \*\* $p < .01$ , \*\*\* $p < .001$ ; CI = Confidence Interval; SE = Standard Error; Z-stat = Z statistic; P-val = P-value; Sig. = Significance level.

#### Accuracy

Paired-samples t-tests revealed that the seen trials had significantly higher ACC than unseen trials (*t*(34) = 16.427, *p* < .001, Cohen’s *d* = 3.418, BF_10_ = 5.318 *×* 10^14^).

The LMMs analysis of ACC revealed that the SOA was not a significant predictor (*χ*^2^(5) = 5.728, *p* = .333; see Figure S4).

**Figure S4.**
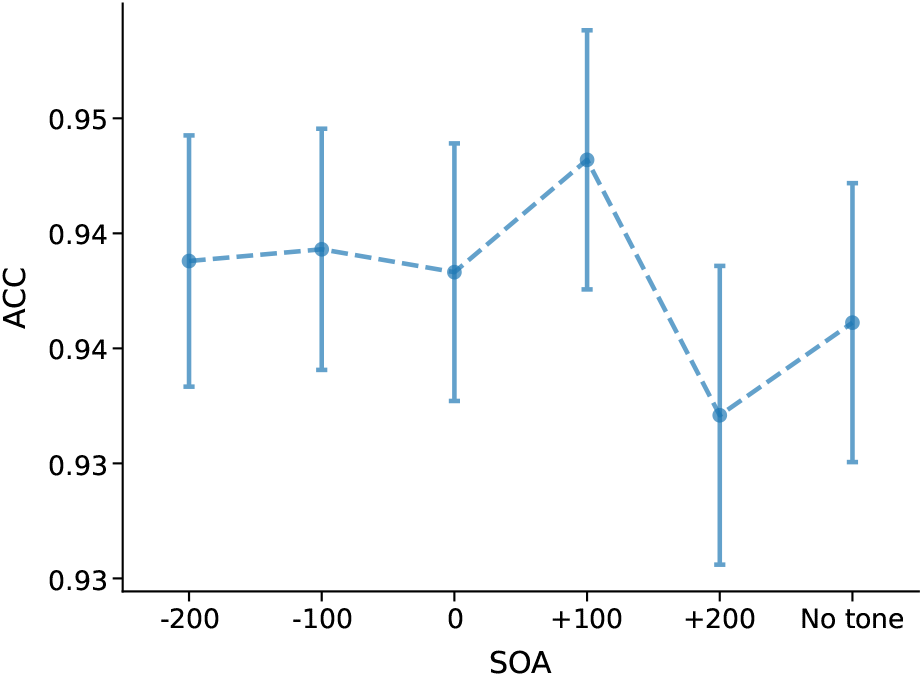
Accuracy across SOA and Awareness conditions in Experiment 2. Error bars represent standard errors.

**Table S6.** Post-hoc pairwise comparisons of ACC for each level of SOA in Experiment 2.

| Contrast | Estimate | CI [2.5%, 97.5%] | SE | Z-stat | P-val | Sig |
| --- | --- | --- | --- | --- | --- | --- |
| (-200) - (-100) | .173 | [-.392, .738] | .238 | .727 | > .05 |  |
| (-200) - (0) | .049 | [-.528, .625] | .242 | .200 | > .05 |  |
| (-200) - (+100) | -.553 | [-1.157, .051] | .255 | -2.167 | > .05 |  |
| (-200) - (+200) | .183 | [-.407, .773] | .248 | .738 | > .05 |  |
| (-200) - (No tone) | -.168 | [-.743, .406] | .241 | -.699 | > .05 |  |
| (-100) - (0) | -.125 | [-.682, .433] | .234 | -.531 | > .05 |  |
| (-100) - (+100) | -.726 | [-1.314, -.138] | .247 | -2.937 | > .05 |  |
| (-100) - (+200) | .010 | [-.559, .578] | .239 | .040 | > .05 |  |
| (-100) - (No tone) | -.341 | [-.901, .218] | .235 | -1.455 | > .05 |  |
| (0) - (+100) | -.601 | [-1.198, -.005] | .251 | -2.396 | > .05 |  |
| (0) - (+200) | .134 | [-.443, .711] | .243 | .552 | > .05 |  |
| (0) - (No tone) | -.216 | [-.783, .351] | .238 | -.909 | > .05 |  |
| (+100) - (+200) | .736 | [.127, 1.344] | .255 | 2.880 | > .05 |  |
| (+100) - (No tone) | .385 | [-.211, .980] | .250 | 1.540 | > .05 |  |
| (+200) - (No tone) | -.351 | [-.916, .214] | .237 | -1.480 | > .05 |  |

Note: \* $p < .05$ , \*\* $p < .01$ , \*\*\* $p < .001$ ; CI = Confidence Interval; SE = Standard Error; Z-stat = Z statistic; P-val = P-value; Sig. = Significance level.

### Proportion seen and SDT tables

**Table S7.**
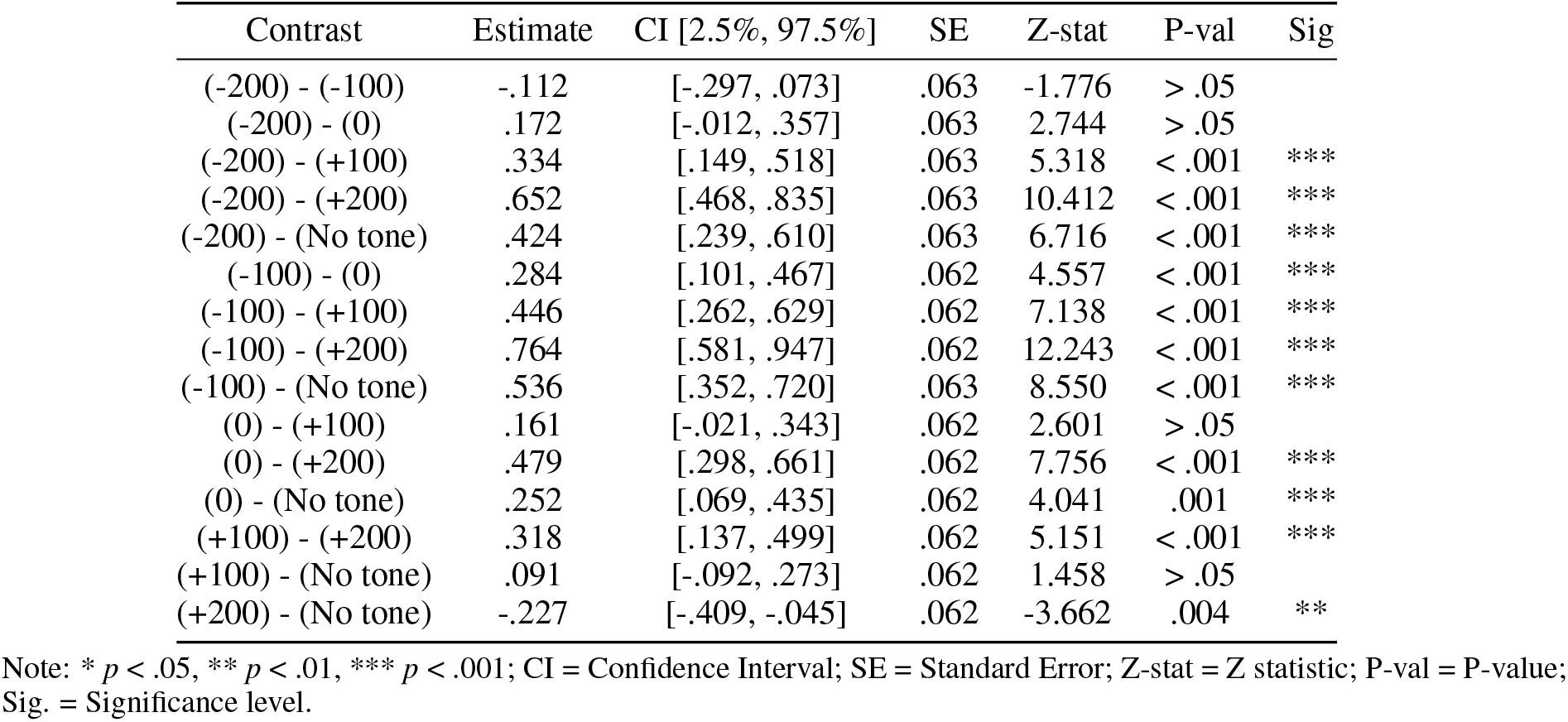
Post-hoc pairwise comparisons of Seen for each level of SOA in Experiment 2.

| Contrast | Estimate | CI [2.5%, 97.5%] | SE | Z-stat | P-val | Sig |
| --- | --- | --- | --- | --- | --- | --- |
| (-200) - (-100) | -.112 | [-.297, .073] | .063 | -1.776 | > .05 |  |
| (-200) - (0) | .172 | [-.012, .357] | .063 | 2.744 | > .05 |  |
| (-200) - (+100) | .334 | [.149, .518] | .063 | 5.318 | < .001 | *** |
| (-200) - (+200) | .652 | [.468, .835] | .063 | 10.412 | < .001 | *** |
| (-200) - (No tone) | .424 | [.239, .610] | .063 | 6.716 | < .001 | *** |
| (-100) - (0) | .284 | [.101, .467] | .062 | 4.557 | < .001 | *** |
| (-100) - (+100) | .446 | [.262, .629] | .062 | 7.138 | < .001 | *** |
| (-100) - (+200) | .764 | [.581, .947] | .062 | 12.243 | < .001 | *** |
| (-100) - (No tone) | .536 | [.352, .720] | .063 | 8.550 | < .001 | *** |
| (0) - (+100) | .161 | [-.021, .343] | .062 | 2.601 | > .05 |  |
| (0) - (+200) | .479 | [.298, .661] | .062 | 7.756 | < .001 | *** |
| (0) - (No tone) | .252 | [.069, .435] | .062 | 4.041 | .001 | *** |
| (+100) - (+200) | .318 | [.137, .499] | .062 | 5.151 | < .001 | *** |
| (+100) - (No tone) | .091 | [-.092, .273] | .062 | 1.458 | > .05 |  |
| (+200) - (No tone) | -.227 | [-.409, -.045] | .062 | -3.662 | .004 | ** |

Note: \* $p < .05$ , \*\* $p < .01$ , \*\*\* $p < .001$ ; CI = Confidence Interval; SE = Standard Error; Z-stat = Z statistic; P-val = P-value; Sig. = Significance level.

**Table S8.** Post-hoc pairwise comparisons of *d*^′^ for each level of SOA in Experiment 2.

| Contrast | T-stat | P-val | BF10 | Cohen's $d$ | Sig. |
| --- | --- | --- | --- | --- | --- |
| (-200) - (-100) | -.753 | > .05 | .236 | -.105 |  |
| (-200) - (0) | .204 | > .05 | .185 | .031 |  |
| (-200) - (+100) | .655 | > .05 | .221 | .116 |  |
| (-200) - (+200) | 2.616 | .197 | 3.385 | .380 |  |
| (-200) - (No tone) | 2.090 | .662 | 1.246 | .357 |  |
| (-100) - (0) | .892 | > .05 | .262 | .146 |  |
| (-100) - (+100) | 1.404 | > .05 | .445 | .236 |  |
| (-100) - (+200) | 3.895 | .007 | 66.329 | .498 | ** |
| (-100) - (No tone) | 3.109 | > .05 | 9.87 | .476 |  |
| (0) - (+100) | .586 | > .05 | .213 | .095 |  |
| (0) - (+200) | 2.785 | > .05 | 4.82 | .390 |  |
| (0) - (No tone) | 2.066 | > .05 | 1.194 | .365 |  |
| (+100) - (+200) | 2.153 | > .05 | 1.392 | .309 |  |
| (+100) - (No tone) | 1.511 | > .05 | .51 | .282 |  |
| (+200) - (No tone) | -.210 | > .05 | .185 | -.034 |  |

Note: \* $p < .05$ , \*\* $p < .01$ , \*\*\* $p < .001$ ; CI = Confidence Interval; SE = Standard Error; Z-stat = Z statistic; P-val = P-value; Sig. = Significance level.

**Table S9.**
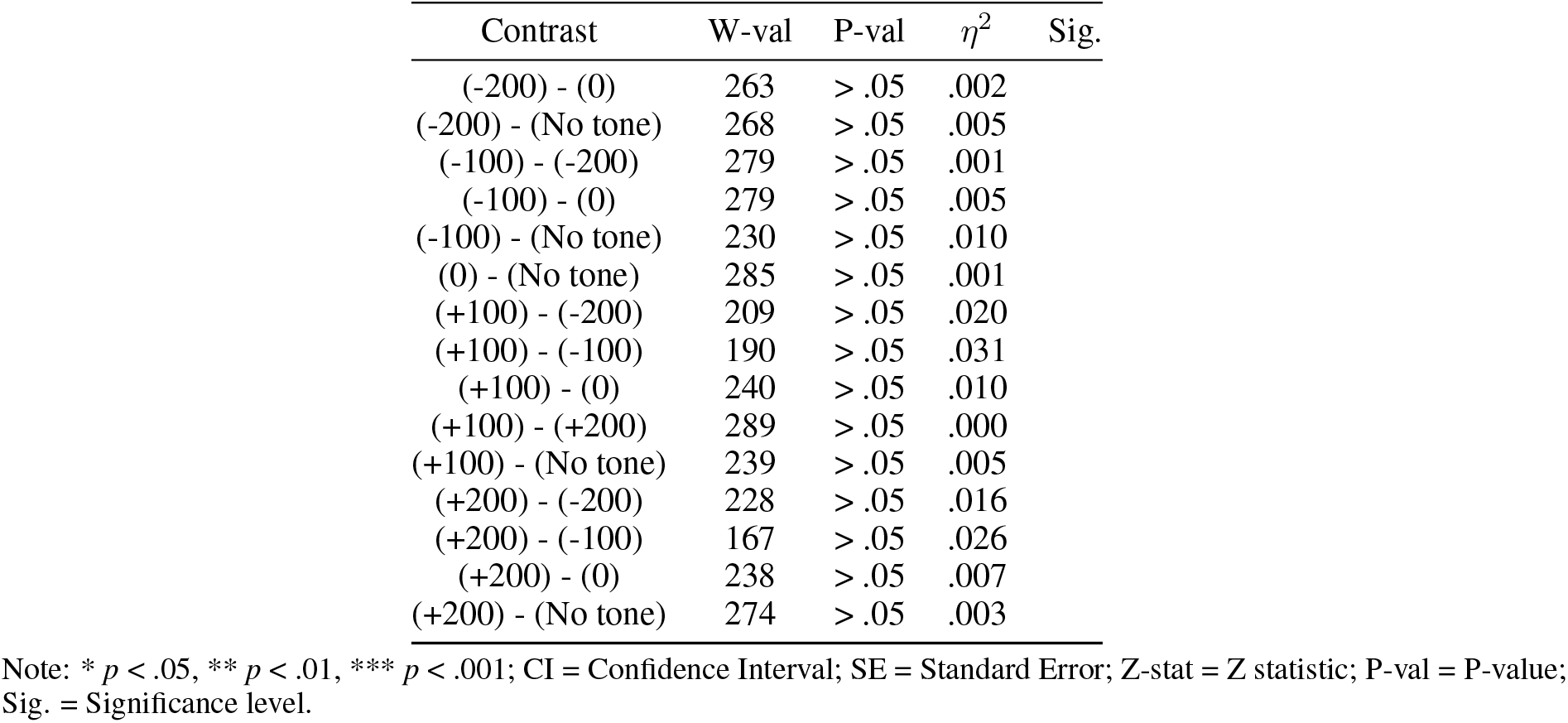
Post-hoc pairwise comparisons of *β* for each level of SOA in Experiment 2.

### Experiment 3

#### Reaction times

T-tests showed significant differences in RTs across Awareness levels, with responses in the seen trials being significantly slower than in the unseen trials (*t*(34) = 14.425, *p* < .001, Cohen’s *d* = 2.831, BF_10_ = 1.272 *×* 10^13^).

**Figure S5.**
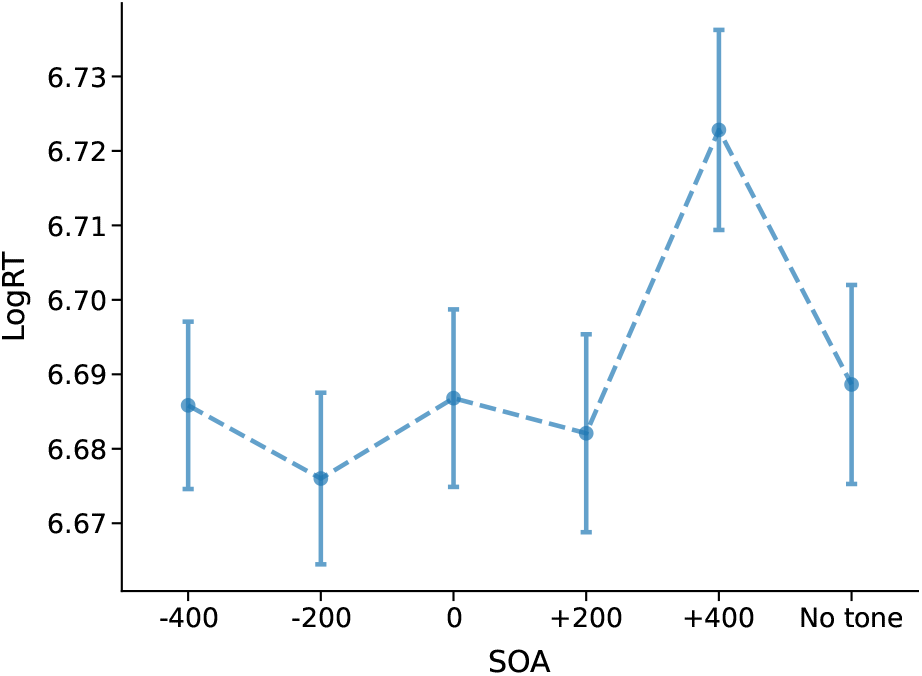
Reaction times (RT) across SOA conditions in Experiment 3. Error bars represent standard errors.

For the LMMs, backwards LRTs revealed a significant effect of SOA (*χ*^2^ (5) = 721.790, *p* < .001), with responses being significantly faster when the tone was presented before or simultaneous to the target onset as shown in Figure S5, although no pairwise comparisons agains the No tone condition were significant (see table S10).

#### Accuracy

The paired-samples t-test showed significant differences in ACC across Awareness levels, with higher accuracy in the seen trials compared to unseen trials (*t*(34) = 30.678, *p* < .001, Cohen’s *d* = 6.089, BF_10_ = 1.16 *×* 10^23^).

In the LMM analysis, the backwards LRTs found no significant effect of SOA (*χ*^2^ (5) = 8.191, *p* > .05).

**Table S10.** Post-hoc pairwise comparisons of RT for each level of SOA in Experiment 3.

| Contrast | Estimate | CI [2.5%, 97.5%] | SE | T-stat | P-val | Sig |
| --- | --- | --- | --- | --- | --- | --- |
| (-400) - (-200) | .010 | [-.028, .047] | .013 | .749 | > .05 |  |
| (-400) - (0) | -.003 | [-.043, .036] | .013 | -.260 | > .05 |  |
| (-400) - (+200) | .011 | [-.031, .052] | .014 | .757 | > .05 |  |
| (-400) - (+400) | -.047 | [-.090, -.005] | .014 | -3.305 | .014 | * |
| (-400) - (No tone) | -.023 | [-.064, .019] | .014 | -1.623 | > .05 |  |
| (-200) - (0) | -.013 | [-.052, .026] | .013 | -.976 | > .05 |  |
| (-200) - (+200) | .001 | [-.040, .043] | .014 | .078 | > .05 |  |
| (-200) - (+400) | -.057 | [-.099, -.015] | .014 | -3.952 | .001 | ** |
| (-200) - (No tone) | -.032 | [-.074, .009] | .014 | -2.292 | > .05 |  |
| (0) - (+200) | .014 | [-.029, .057] | .015 | .964 | > .05 |  |
| (0) - (+400) | -.044 | [-.088, -.000] | .015 | -2.951 | .048 | * |
| (0) - (No tone) | -.019 | [-.062, .024] | .015 | -1.323 | > .05 |  |
| (+200) - (+400) | -.058 | [-.104, -.012] | .016 | -3.730 | .003 | ** |
| (+200) - (No tone) | -.034 | [-.079, .011] | .015 | -2.190 | > .05 |  |
| (+400) - (No tone) | .025 | [-.021, .070] | .016 | 1.581 | > .05 |  |

Note: \* $p < .05$ , \*\* $p < .01$ , \*\*\* $p < .001$ ; CI = Confidence Interval; SE = Standard Error; Z-stat = Z statistic; P-val = P-value; Sig. = Significance level.

**Figure S6.**
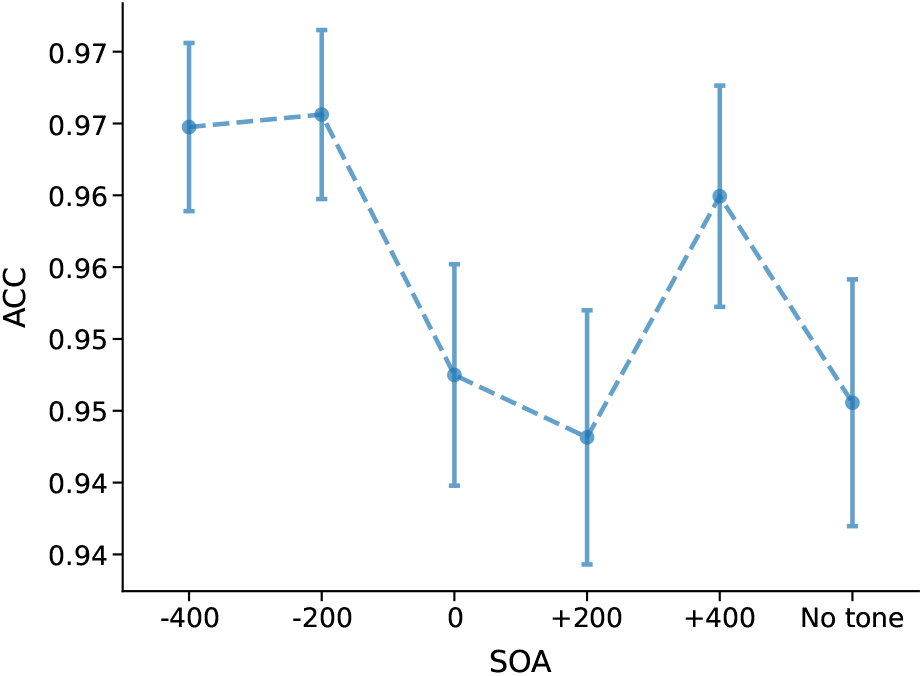
ACC across SOA conditions in Experiment 3. Error bars represent standard errors.

**Table S11.** Post-hoc pairwise comparisons of ACC for each level of SOA in Experiment 3.

| Contrast | Estimate | CI [2.5%, 97.5%] | SE | Z-stat | P-val | Sig |
| --- | --- | --- | --- | --- | --- | --- |
| (-400) - (-200) | -.034 | [-.768, .700] | .250 | -.137 | > .05 |  |
| (-400) - (0) | .319 | [-.376, 1.014] | .237 | 1.346 | > .05 |  |
| (-400) - (+200) | .452 | [-.266, 1.170] | .245 | 1.847 | > .05 |  |
| (-400) - (+400) | -.052 | [-.842, .739] | .269 | -.192 | > .05 |  |
| (-400) - (No tone) | .396 | [-.326, 1.118] | .246 | 1.611 | > .05 |  |
| (-200) - (0) | .353 | [-.351, 1.056] | .240 | 1.472 | > .05 |  |
| (-200) - (+200) | .486 | [-.242, 1.214] | .248 | 1.961 | > .05 |  |
| (-200) - (+400) | -.018 | [-.817, .782] | .272 | -.065 | > .05 |  |
| (-200) - (No tone) | .430 | [-.300, 1.161] | .249 | 1.729 | > .05 |  |
| (0) - (+200) | .133 | [-.554, .821] | .234 | .570 | > .05 |  |
| (0) - (+400) | -.370 | [-1.132, .391] | .260 | -1.427 | > .05 |  |
| (0) - (No tone) | .078 | [-.613, .768] | .235 | .330 | > .05 |  |
| (+200) - (+400) | -.504 | [-1.287, .279] | .267 | -1.889 | > .05 |  |
| (+200) - (No tone) | -.056 | [-.768, .657] | .243 | -.230 | > .05 |  |
| (+400) - (No tone) | .448 | [-.338, 1.234] | .268 | 1.674 | > .05 |  |

Note: \* $p < .05$ , \*\* $p < .01$ , \*\*\* $p < .001$ ; CI = Confidence Interval; SE = Standard Error; Z-stat = Z statistic; P-val = P-value; Sig. = Significance level.

### Proportion seen and SDT tables

**Table S12.**
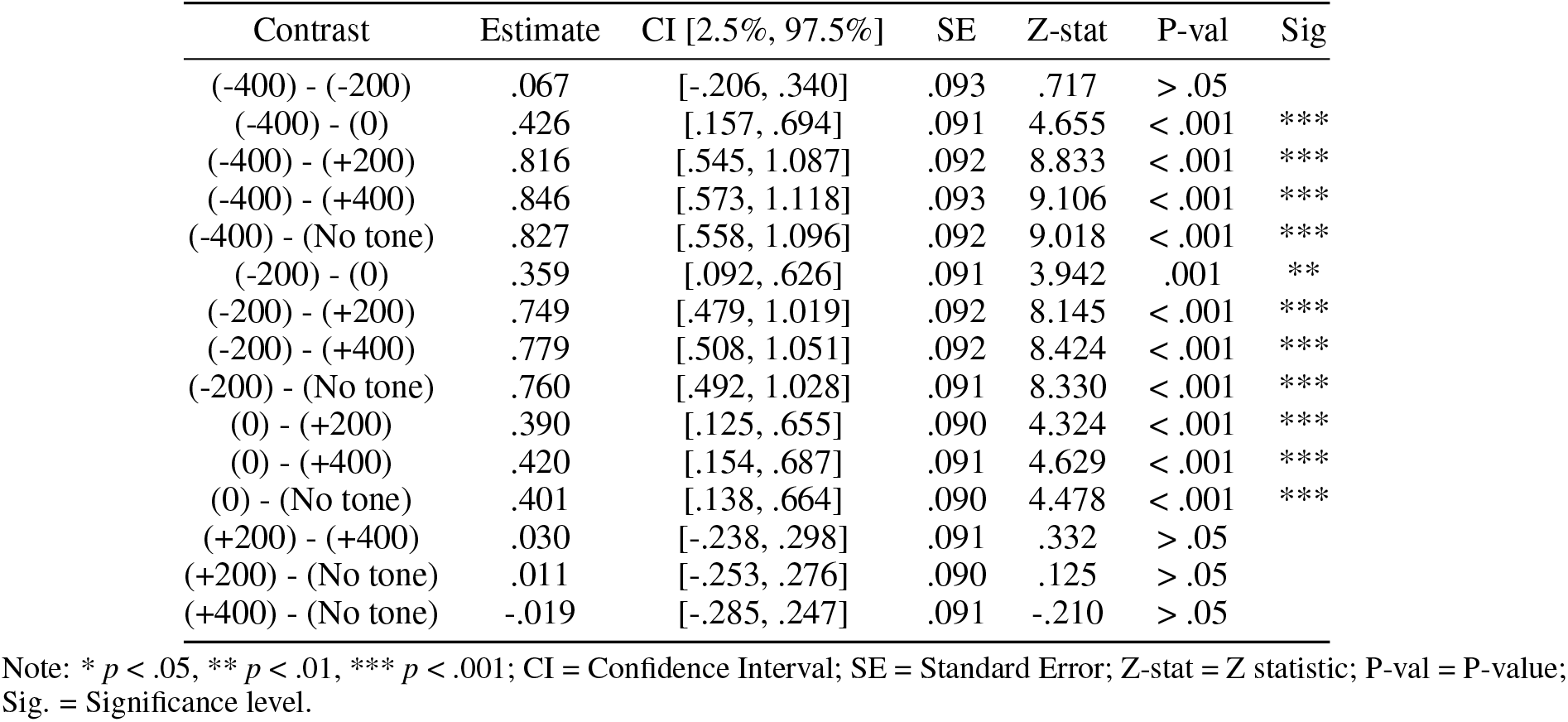
Post-hoc pairwise comparisons of Seen for each level of SOA in Experiment 3.

**Table S13.** Post-hoc pairwise comparisons of *d*^′^ for each level of SOA in Experiment 3.

| Contrast | W-val | P-val | $\eta^2$ | Sig. |
| --- | --- | --- | --- | --- |
| (-400) - (0) | 152.000 | .100 | .040 |  |
| (-400) - (No tone) | 32.000 | .000 | .160 | *** |
| (-200) - (-400) | 287.000 | 1.000 | .001 |  |
| (-200) - (0) | 132.000 | .209 | .028 |  |
| (-200) - (No tone) | 32.000 | .000 | .142 | *** |
| (0) - (No tone) | 88.000 | .001 | .078 | ** |
| (+200) - (-400) | 64.000 | .000 | .091 | *** |
| (+200) - (-200) | 70.000 | .000 | .075 | *** |
| (+200) - (0) | 209.000 | 1.000 | .022 |  |
| (+200) - (+400) | 275.000 | 1.000 | .000 |  |
| (+200) - (No tone) | 190.000 | .606 | .018 |  |
| (+400) - (-400) | 70.000 | .000 | .107 | *** |
| (+400) - (-200) | 99.000 | .003 | .089 | ** |
| (+400) - (0) | 180.000 | .395 | .028 |  |
| (+400) - (No tone) | 219.000 | 1.000 | .021 |  |

Note: \* $p < .05$ , \*\* $p < .01$ , \*\*\* $p < .001$ ; CI = Confidence Interval; SE = Standard Error; Z-stat = Z statistic; P-val = P-value; Sig. = Significance level.

**Table S14.** Post-hoc pairwise comparisons of *β* for each level of SOA in Experiment 3.

| Contrast | W-val | P-val | $\eta^2$ | Sig. |
| --- | --- | --- | --- | --- |
| (-400) - (0) | 253.000 | 1.000 | .001 |  |
| (-400) - (No tone) | 204.000 | 1.000 | .014 |  |
| (-200) - (-400) | 249.000 | 1.000 | .007 |  |
| (-200) - (0) | 247.000 | 1.000 | .002 |  |
| (-200) - (No tone) | 172.000 | .274 | .038 |  |
| (0) - (No tone) | 193.000 | .685 | .024 |  |
| (+200) - (-400) | 258.000 | 1.000 | .007 |  |
| (+200) - (-200) | 289.000 | 1.000 | .000 |  |
| (+200) - (0) | 304.000 | 1.000 | .002 |  |
| (+200) - (+400) | 290.000 | 1.000 | .000 |  |
| (+200) - (No tone) | 140.000 | .051 | .042 |  |
| (+400) - (-400) | 260.000 | 1.000 | .011 |  |
| (+400) - (-200) | 309.000 | 1.000 | .000 |  |
| (+400) - (0) | 309.000 | 1.000 | .004 |  |
| (+400) - (No tone) | 157.000 | .130 | .054 |  |

Note: \* $p < .05$ , \*\* $p < .01$ , \*\*\* $p < .001$ ; CI = Confidence Interval; SE = Standard Error; Z-stat = Z statistic; P-val = P-value; Sig. = Significance level.

### Experiment 4

#### Reaction times

First, we compared RTs for seen versus unseen trials by means of a t-test, which revealed significantly faster RTs for seen trials compared to unseen trials (*t*(34) = 9.772, *p* < .001, Cohen’s *d* = 1.526, BF_10_ = 4.476 *×* 10^8^).

For the LMMs, backwards LRTs revealed a significant main effect of SOA (*χ*^2^ (5) = 39.615, *p* < .001), with faster responses when the tone was presented before or simultaneously with the target, compared to when the tone was presented after the target (see Figure S7).

**Figure S7.**
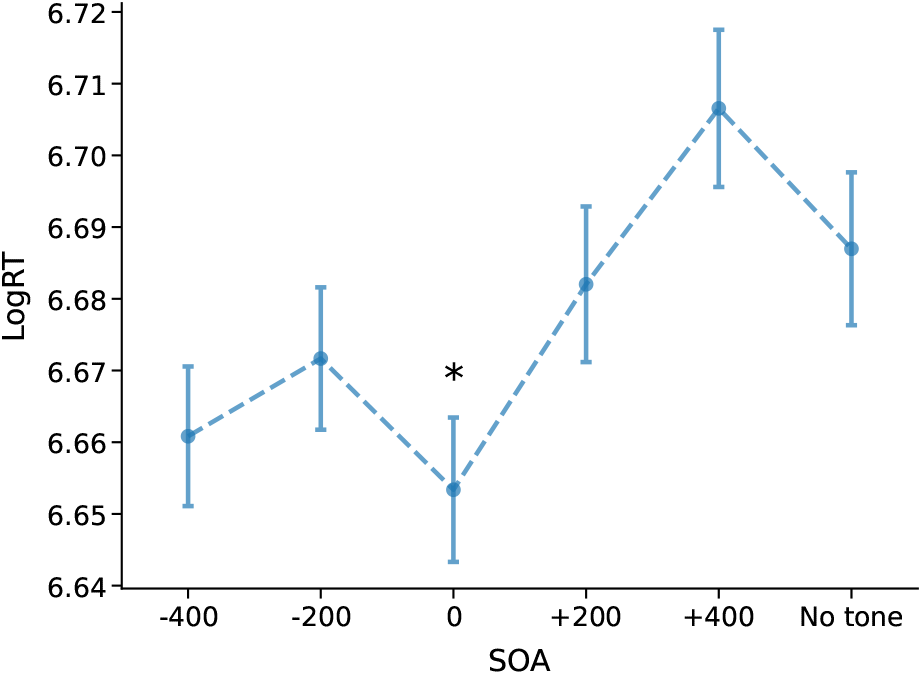
RTs across SOA conditions in Experiment 4. Error bars represent standard errors. Asterisks indicate significant post-hoc comparisons against the No tone condition

**Table S15.** Post-hoc pairwise comparisons of RT for each level of SOA in Experiment 4.

| Contrast | Estimate | CI [2.5%, 97.5%] | SE | T-stat | P-val | Sig |
| --- | --- | --- | --- | --- | --- | --- |
| (-400) - (-200) | -.007 | [-.037, .023] | .010 | -.723 | > .05 |  |
| (-400) - (0) | .010 | [-.021, .041] | .011 | .950 | > .05 |  |
| (-400) - (+200) | -.026 | [-.058, .006] | .011 | -2.403 | > .05 |  |
| (-400) - (+400) | -.052 | [-.084, -.020] | .011 | -4.716 | < .001 | *** |
| (-400) - (No tone) | -.024 | [-.057, .008] | .011 | -2.220 | > .05 |  |
| (-200) - (0) | .017 | [-.013, .048] | .010 | 1.676 | > .05 |  |
| (-200) - (+200) | -.019 | [-.050, .013] | .011 | -1.749 | > .05 |  |
| (-200) - (+400) | -.045 | [-.077, -.013] | .011 | -4.091 | .001 | *** |
| (-200) - (No tone) | -.017 | [-.049, .015] | .011 | -1.565 | > .05 |  |
| (0) - (+200) | -.036 | [-.069, -.004] | .011 | -3.281 | .016 | * |
| (0) - (+400) | -.062 | [-.095, -.029] | .011 | -5.554 | < .001 | *** |
| (0) - (No tone) | -.034 | [-.067, -.002] | .011 | -3.092 | .030 | * |
| (+200) - (+400) | -.026 | [-.060, .008] | .012 | -2.239 | > .05 |  |
| (+200) - (No tone) | .002 | [-.032, .036] | .011 | .161 | > .05 |  |
| (+400) - (No tone) | .028 | [-.006, .062] | .012 | 2.385 | > .05 |  |

Note: \* $p < .05$ , \*\* $p < .01$ , \*\*\* $p < .001$ ; CI = Confidence Interval; SE = Standard Error; Z-stat = Z statistic; P-val = P-value; Sig. = Significance level.

#### Accuracy

Paired-samples t-tests comparing accuracy between seen and unseen trials revealed significantly higher accuracy for seen trials compared to unseen trials (*t*(34) = 23.786, *p* < .001, Cohen’s *d* = 5.130, BF_10_ = 3.852 *×* 10^19^).

For the LMM analysis, backwards LRTs revealed a significant main effect of SOA (*χ*^2^ (5) = 12.220, *p* = .031). Post-hoc pairwise comparisons indicated a general effect of the tone increasing the ACC when compared to no presenting the tone, regardless of the SOA (see Figure S8 and Table S16).

**Figure S8.**
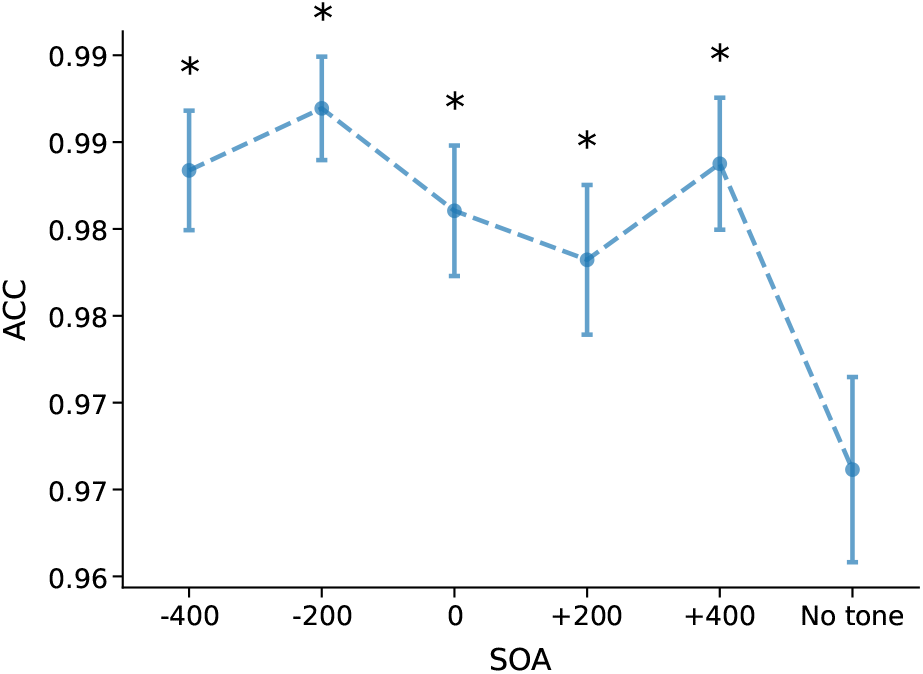
ACC across SOA conditions in Experiment 4. Error bars represent standard errors. Asterisks indicate significant post-hoc comparisons against the No tone condition.

**Table S16.** Post-hoc pairwise comparisons of ACC for each level of SOA in Experiment 4.

| Contrast | Estimate | CI [2.5%, 97.5%] | SE | Z-stat | P-val | Sig |
| --- | --- | --- | --- | --- | --- | --- |
| (-400) - (-200) | -.276 | [-.279, -.274] | .001 | -324.098 | < .001 | *** |
| (-400) - (0) | .221 | [.219, .224] | .001 | 259.226 | < .001 | *** |
| (-400) - (+200) | .209 | [.207, .212] | .001 | 245.547 | < .001 | *** |
| (-400) - (+400) | -.105 | [-.108, -.103] | .001 | -123.446 | < .001 | *** |
| (-400) - (No tone) | .706 | [.703, .708] | .001 | 826.984 | < .001 | *** |
| (-200) - (0) | .498 | [.494, .501] | .001 | 412.480 | < .001 | *** |
| (-200) - (+200) | .486 | [.482, .490] | .001 | 402.808 | < .001 | *** |
| (-200) - (+400) | .171 | [.168, .175] | .001 | 141.885 | < .001 | *** |
| (-200) - (No tone) | .982 | [.978, .986] | .001 | 813.954 | < .001 | *** |
| (0) - (+200) | -.012 | [-.015, -.008] | .001 | -9.681 | < .001 | *** |
| (0) - (+400) | -.326 | [-.330, -.323] | .001 | -270.597 | < .001 | *** |
| (0) - (No tone) | .484 | [.481, .488] | .001 | 401.456 | < .001 | *** |
| (+200) - (+400) | -.315 | [-.318, -.311] | .001 | -260.922 | < .001 | *** |
| (+200) - (No tone) | .496 | [.492, .500] | .001 | 411.146 | < .001 | *** |
| (+400) - (No tone) | .811 | [.807, .814] | .001 | 672.068 | < .001 | *** |

Note: \* $p < .05$ , \*\* $p < .01$ , \*\*\* $p < .001$ ; CI = Confidence Interval; SE = Standard Error; Z-stat = Z statistic; P-val = P-value; Sig. = Significance level.

### Proportion seen and SDT tables

**Table S17.** Post-hoc pairwise comparisons of Seen for each level of SOA in Experiment 4.

| Contrast | Estimate | CI [2.5%, 97.5%] | SE | Z-stat | P-val | Sig |
| --- | --- | --- | --- | --- | --- | --- |
| (-400) - (-200) | -.138 | [-.522, .246] | .131 | -1.054 | > .05 |  |
| (-400) - (0) | .378 | [.022, .734] | .121 | 3.120 | .027 | * |
| (-400) - (+200) | .721 | [.375, 1.068] | .118 | 6.112 | < .001 | *** |
| (-400) - (+400) | .981 | [.646, 1.317] | .114 | 8.588 | < .001 | *** |
| (-400) - (No tone) | 1.027 | [.694, 1.359] | .113 | 9.066 | < .001 | *** |
| (-200) - (0) | .516 | [.154, .877] | .123 | 4.190 | < .001 | *** |
| (-200) - (+200) | .859 | [.507, 1.212] | .120 | 7.159 | < .001 | *** |
| (-200) - (+400) | 1.119 | [.778, 1.461] | .116 | 9.614 | < .001 | *** |
| (-200) - (No tone) | 1.165 | [.826, 1.503] | .115 | 10.094 | < .001 | *** |
| (0) - (+200) | .343 | [.023, .664] | .109 | 3.142 | .025 | * |
| (0) - (+400) | .604 | [.295, .913] | .105 | 5.733 | < .001 | *** |
| (0) - (No tone) | .649 | [.343, .954] | .104 | 6.231 | < .001 | *** |
| (+200) - (+400) | .260 | [-.038, .558] | .102 | 2.563 | > .05 |  |
| (+200) - (No tone) | .305 | [.011, .600] | .100 | 3.043 | .035 | * |
| (+400) - (No tone) | .045 | [-.236, .327] | .096 | .471 | > .05 |  |

Note: \* $p < .05$ , \*\* $p < .01$ , \*\*\* $p < .001$ ; CI = Confidence Interval; SE = Standard Error; Z-stat = Z statistic; P-val = P-value; Sig. = Significance level.

**Table S18.** Post-hoc pairwise comparisons of *d*^′^ for each level of SOA in Experiment 4.

| Contrast | T-stat | P-val | BF10 | Cohen's $d$ | Sig. |
| --- | --- | --- | --- | --- | --- |
| (-400) - (-200) | -.491 | >.05 | .203 | -.090 |  |
| (-400) - (0) | 1.120 | >.05 | .323 | .249 |  |
| (-400) - (+200) | 2.663 | >.05 | 3.729 | .463 |  |
| (-400) - (+400) | 3.229 | .041 | 13.008 | .640 | * |
| (-400) - (No tone) | 4.730 | .001 | 601.236 | .900 | *** |
| (-200) - (0) | 2.258 | >.05 | 1.684 | .348 |  |
| (-200) - (+200) | 3.299 | .034 | 15.334 | .588 | * |
| (-200) - (+400) | 3.532 | .018 | 26.834 | .763 | * |
| (-200) - (No tone) | 5.699 | .000 | 8628.595 | 1.042 | *** |
| (0) - (+200) | .842 | >.05 | .252 | .179 |  |
| (0) - (+400) | 2.021 | >.05 | 1.106 | .375 |  |
| (0) - (No tone) | 3.170 | .048 | 11.339 | .626 | * |
| (+200) - (+400) | 1.121 | >.05 | .323 | .244 |  |
| (+200) - (No tone) | 3.127 | >.05 | 10.27 | .541 |  |
| (+400) - (No tone) | 1.420 | >.05 | .454 | .258 |  |

Note: \* $p < .05$ , \*\* $p < .01$ , \*\*\* $p < .001$ ; CI = Confidence Interval; SE = Standard Error; Z-stat = Z statistic; P-val = P-value; Sig. = Significance level.

**Table S19.**
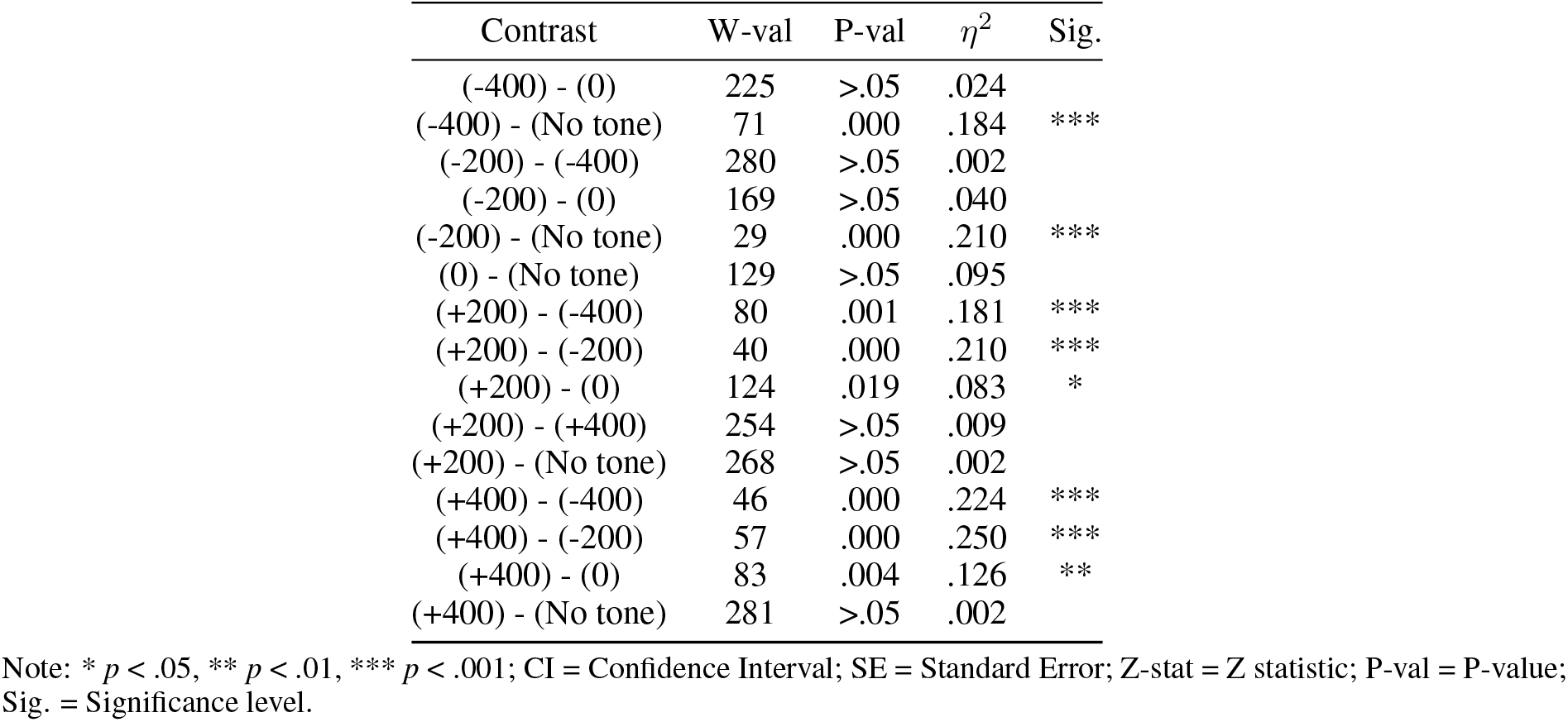
Post-hoc pairwise comparisons of *β* for each level of SOA in Experiment 4.

### Comparison between Experiment 3 and Experiment 4

To more directly assess whether the retro-cue effect depended on target visibility, an additional between-experiments comparison was conducted including Experiments 3 and 4, which used the same SOAs but differed in target visibility (50% vs. 75%). Specifically, a combined mixed-effects model was fitted with Experiment and SOA as fixed factors:

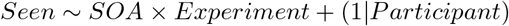

This analysis revealed a strong main effect of Experiment, with higher overall seen rates in Experiment 4 (.857) than in Experiment 3 (.644), consistent with the higher visibility range introduced by the titration procedure. Importantly, there was also a small but significant SOA × Experiment interaction (*χ*^2^(5) = 11.36, *p* = .045), indicating that the modulation of seen responses across SOAs differed between the two experiments.

To further examine whether the retro-cue benefit itself differed across experiments, an index of retro-cue benefit was computed for each SOA by subtracting the proportion of seen responses in the No tone condition from the proportion of seen responses in each SOA condition. This index was then directly compared between Experiments 3 and 4. In this analysis, the SOA × Experiment interaction did not reach significance (*p* = .060). A planned comparison at the +200 ms SOA showed a numerically larger retro-cue benefit in Experiment 4 than in Experiment 3, but this difference did not reach significance (*p* = .235). Table S20 reports the pairwise contrasts (Experiment 3 vs Experiment 4) for each SOA, including confidence intervals and test statistics.

Taken together, these supplementary analyses provide converging but not definitive support for the visibility-dependent interpretation of the retro-cue effect. Because this between-experiments comparison was not planned *a priori* and the study was not powered specifically for this contrast, these results should be interpreted cautiously as trend-level support rather than as conclusive evidence.

**Table S20.**
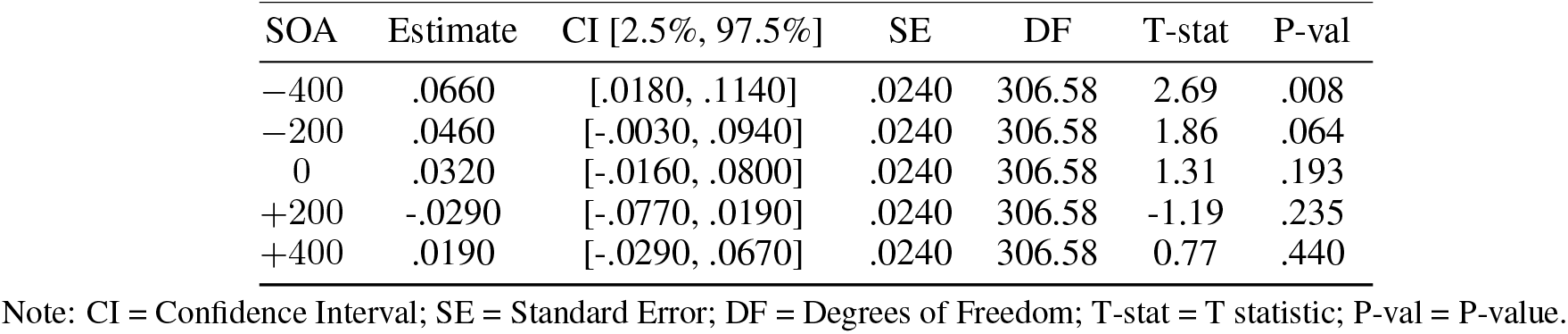
Pairwise contrasts for the comparison between Experiment 3 and Experiment 4 across SOAs, in the proportion of seen index.

